# A moonlighting function of a chitin polysaccharide monooxygenase, CWR-1, in allorecognition in *Neurospora crassa*

**DOI:** 10.1101/2022.06.26.497659

**Authors:** Tyler C. Detomasi, Adriana M. Rico-Ramírez, Richard I. Sayler, A. Pedro Gonçalves, Michael A. Marletta, N. Louise Glass

**Affiliations:** Department of Chemistry, University of California, Berkeley, Berkeley, CA 94720, USA; Department of Plant and Microbial Biology, University of California, Berkeley, Berkeley, CA 94720, USA; California Institute for Quantitative Biosciences, University of California, Berkeley, Berkeley, CA 94720, USA; Department of Molecular and Cell Biology, University of California, Berkeley, Berkeley, CA 94720, USA; Department of Cell Biology and Anatomy, College of Medicine, National Cheng Kung University, Tainan City, 701, Taiwan

## Abstract

Organisms require the ability to differentiate themselves from organisms of different or even the same species. Allorecognition processes in filamentous fungi are essential to ensure identity of an interconnected syncytial colony to protect it from exploitation and disease. *Neurospora crassa* has three cell fusion checkpoints controlling formation of an interconnected mycelial network. The locus that controls the second checkpoint, which allows for cell wall dissolution and subsequent fusion between cells/hyphae, *cwr,* encodes two linked genes, *cwr-1* and *cwr-2*. Previously, it was shown that *cwr-1* and *cwr-2* show severe linkage disequilibrium with six different haplogroups present in *N. crassa* populations. Isolates from an identical *cwr* haplogroup show robust fusion, while somatic cell fusion between isolates of different haplogroups is significantly blocked in cell wall dissolution. The *cwr-1* gene encodes a putative polysaccharide monooxygenase (PMO). Herein we confirm that CWR-1 is a C1-oxidizing chitin PMO. We show that the PMO domain of CWR-1 was sufficient for checkpoint function and cell fusion blockage; however, through analysis of active-site, histidine-brace mutants, the catalytic activity of CWR-1 was ruled out as a major factor for allorecognition. Swapping a portion of the PMO domain (V86 to T130) did not switch *cwr* haplogroup specificity, but rather cells containing this chimera exhibited a novel haplogroup specificity. Allorecognition to mediate cell fusion blockage is likely occurring through a protein-protein interaction between CWR-1 with CWR-2. These data highlight a moonlighting role in allorecognition of the CWR1 PMO domain.

## Introduction

Allorecognition is the ability of a cell to recognize self or kin and has widespread importance for many organisms that have multicellular organization. Allorecognition in some form is utilized by the social amoeba, *Dictyostelium discoideum* (Kundert and Shaulsky, 2019; Kuzdzal-Fick, et al., 2011) the bacterium, *Proteus mirabilis* (Gibbs, Mark, and Greenberg, 2008; Gibbs and Greenberg, 2011), gymnosperms (Pandey, 1960), slime molds (Clark, 2003; Shaulsky and Kessin, 2007), and invertebrates, such as *Botryllus schlosseri* (De Tomaso et al., 2005; Rosengarten and Nicotra, 2011; Yoshito et al., 2008) or the cnidarian *Hydractinia symbiolongicarpus* (Rosengarten and Nicotra, 2011). In *Drosophila melanogaster,* the *dscam* gene, which contains >19,000 splicing isoforms, uses differential splicing and homo-dimer binding to recognize and not synapse with itself (Wojtowicz, et al. 2004; Wojtowicz et al., 2007). In vertebrate species, the major histocompatibility complex (MHC) is crucial for the immune response and allows cells to identify infected cells or cells that are no longer kin for destruction (Afzali et al. 2008; Marino et al., 2016).

Although not strictly multicellular, most filamentous fungi grow as an interconnected hyphal network that shares cytoplasm, nuclei, and nutrients and utilizes allorecognition machinery to regulate this process (Gonçalves and Glass, 2020). This syncytial organization of filamentous fungi allows an interconnected colony to thrive in an environment that is heterogeneous for nutrients and habitats (Anna et al. 2012). The formation of an interconnected mycelial network is a process that is controlled by communication and cell fusion checkpoints to help ensure that cells with higher genetic identity form a colony (Gonçalves and Glass, 2020; Gonçalves et al., 2020). Without checkpoints, cell fusion can occur between genetically unrelated colonies and cells, allowing “cheater” nuclei to steal resources (Bastiaans, et al., 2016; Grum-Grzhimaylo et al., 2021) or enabling the spread of cytoplasmic diseases such as mycoviruses(Zhang and Nuss, 2016) or selfish genetic elements (Debets et al., 2012), which are spread via cell fusion throughout fungal populations.

The filamentous ascomycete species, *Neurospora crassa,* has three cell fusion checkpoints that function in allorecognition. The first checkpoint regulates chemotropic interactions between hyphae and/or germinated asexual spores (germlings) and is controlled by the determinant of communication (*doc*) locus that senses chemosignalling (Heller et al., 2016). Cells with identical allelic specificity at the *doc* locus undergo chemotropic interactions, while cells with alternate *doc* allelic specificity show greatly reduced interactions. The second checkpoint is regulated by the cell wall remodeling locus*, cwr*, which contains the two linked genes, *cwr-1* (Uniprot KB Q1K703) and *cwr-2* (UniprotKB Q1K701) (Gonçalves et al., 2019). Cells/hyphae with identical allelic specificity at *doc* and *cwr* undergo chemotropic interactions and, upon contact, undergo cell wall dissolution, membrane merger, and cytoplasmic mixing. However, if cells have identical *doc* specificity, but differ in *cwr* allelic specificity, hyphae/cells undergo chemotropic interactions, but upon contact, cells remain adhered and fail to undergo cell wall deconstruction and membrane merger at the point of contact (Gonçalves et al., 2019). The final checkpoint is a post-fusion checkpoint. Following chemotrophic interactions (*doc* alleles with identical allelic specificity) and cell fusion (*cwr* alleles with identical allelic specificity), if cells/hyphae differ in specificity at several post-fusion allorecognition loci, the fusion cell is compartmentalized and undergoes rapid cell death (Daskalov et al., 2019; Heller et al.,2018; Rico-Ramírez et al., 2022). Recent data in *N. crassa* showed that one of these post-fusion allorecognition loci has functional and structural similarity to mammalian gasdermin and confers rapid cell death via a pyroptotic-like mechanism (Corinne et al., 2022; Daskalov and Glass, 2022; Daskalov et al., 2020; Rico-Ramírez et al., 2022)while in the related filamentous fungus, *Podospora anserina*, a different post-fusion cell death allorecognition locus has functional similarities to necroptosis (Saupe, 2011, 2020).

The *cwr-1* gene is predicted to encode a chitin-active polysaccharide monooxygenase (PMO) in the CAZy (Carbohydrate-Active enZYme) database designation of Auxiliary Activity 11 (AA11) family (Levasseur et al., 2013). The *cwr-2* gene encodes a predicted protein that contains eight transmembrane regions with two annotated domains of unknown function (PF11915). Within populations of *N. crassa*, *cwr-1/cwr-2* alleles show severe linkage disequilibrium and fall into six different haplogroups (HG). Cells/germlings containing only CWR-1 from HG1 are blocked in cell fusion with isogenic cells containing only CWR-2 from a different haplogroup (Levasseur et al., 2013). These data indicate that incompatible CWR-1-CWR-2 function in *trans* when present in different interacting cells and are necessary and sufficient to trigger the cell fusion block.

PMOs, alternatively referred to as lytic polysaccharide monooxygenases (LPMOs), have been studied extensively for over a decade (Phillips et al., 2011; Vaaje-Kolstad et al., 2010). These proteins were first discovered as auxiliary redox enzymes that greatly enhance cellulose degradation in combination with glycosyl hydrolases. Beyond just cellulolytic activity, other PMOs have been demonstrated to show activity on other polysaccharides such as chitin (Hemsworth et al., 2014), starch (Vu et al., 2019; Vu and Marletta, 2016), xylans (Couturier et al., 2018; Hüttner et al., 2019), and various hemicelluloses (Agger et al., 2014; Monclaro et al., 2020). PMOs have been of fundamental interest because they catalyze the hydroxylation of a strong C-H bond and their utility in industrial biofuel applications (Johansen, 2016; Moreau et al., 2019). However, as more PMO families have been identified, other roles have been noted, including in the life cycle of insect viruses (Chiu et al., 2015), insect development (Sabbadin et al., 2018), fungal development (Fu et al.,2014; Gonçalves et al., 2019; Maddi et al., 2012), and in symbiotic associations between bacteria and arthropods (Distel et al., 2011; Pinheiro et al., 2015). All PMOs contain a secretion signal peptide that is cleaved leaving the mature protein with an N-terminal histidine residue that binds a single copper atom (Phillips et al., 2011). Of the four other proteins characterized in the AA11 family, only a chitin-degradation role has been demonstrated (Hemsworth et al., 2014; Rieder et al., 2021; Støpamo et al., 2021; Wang, Li, Salazar-Alvarez, et al., 2018; Wang et al., 2018). Three of the four proteins have a similar protein architecture composed of an N-terminal PMO domain followed by a G/S rich disordered linker and an X278 domain similar in sequence to carbohydrate-binding modules (CBMs) at the C-terminus. If this is a true CBM, it likely binds chitin, as it is also found in GH-18 chitinases (Hemsworth et al., 2014), but no definitive biochemical characterization has been performed.

In this study, we assessed whether CWR-1 from each of the six different haplogroups has PMO activity and defined the substrate and products of that PMO activity. We also assessed whether CWR-1 proteins from each haplogroup formed unique products that would explain CWR-1 allelic specificity. By the further construction of PMO domain chimeras, we evaluated whether a polymorphic region of the PMO domain (LS and LC loop) was a factor in triggering allorecognition in interactions with CWR-2. Our data indicates that CWR-1 has chitin PMO activity, but that catalytic activity was not required for cell fusion blockage, highlighting an evolved and additional function of this PMO in allorecognition.

## Materials and Methods

### Recombinant DNA techniques and plasmid constructs

Plasmids and oligonucleotides used in this work are listed in Appendix 1-Tables 1-3. All plasmids used to transform *N. crassa* are derived from plasmid pMF272 (Freitag et al., 2004) PCR reactions were performed in MiniAmp Plus Thermal Cycler with Q5^®^ High-Fidelity DNA polymerase (2000 U/mL) (NEB Ipswich, Massachusetts) according to the manufacturer’s instructions. Genes were amplified using genomic DNA of *N. crassa* strains (FGSC2489 or P4471, JW228, JW242, JW258, P4476, D111) as a template. The fragments amplified were purified using Monarch^®^ DNA Gel extraction Kit (T1020S NEB, Ipswich, Massachusetts). The *cwr-1* chimeras were made by fusing two fragments, a fragment that encodes the signal peptide and the PMO domain corresponding to alleles from each of the six haplogroups of *cwr-1* (JW228, JW242, JW258, P4471, P4476, D111) and a second fragment that included the region corresponding to the GS linker and X278 domain of *cwr-1^FGSC2489^* (haplogroup 1). The fragments were fused using polymerase Phusion high-fidelity DNA Polymerase (M0530S, NEB, Ipswich, Massachusetts) following the manufacturer’s instructions. We designed primers flanking the ORF of the *cwr-1* (NCU01380), *cwr-2* (NCU01382), and chimeras of *cwr-1* with *Xba*I and *Pac*I restriction sites for the 5’ and 3’ ends, respectively. Amplified and gel-purified PCR ORFs were *Xba*I (R0145S, NEB, Ipswich, Massachusetts) and *Pac*I (R0547S, NEB, Ipswich, Massachusetts) digested and inserted into *Xba*I/*Pac*I digested *pMF272::his-3-*P*tef*-1-V5-T*ccg-1.* The fragments were gel purified and ligated using T4 DNA ligase (15224-017 Invitrogen life technologies, Waltham, Massachusetts). The resulting constructs were used to transform *N. crassa (△NCU01380△NCU01381△NCU01382::hph; his-3^-^)*.

The plasmids to transform *N. crassa* with the truncated version of *cwr-1 (cwr-1△GS^HG1^, cwr-1△GS△X278^HG1^),* histidine mutants *(cwr-1H20A^HG1^, cwr-1H78A^HG1^, cwr-1H20AH78A^HG1^)* and the loop chimera *(cwr-1^HG1^LS^HG6(V86-T130))^* was constructed using the plasmid *pMF272::his-3-Pcwr-1-cwr-1(NCU01380)-Tcwr-1^FGSC2489HG1^*as a backbone. According to the manufacturer’s instructions, the truncated versions of *cwr-1* and the histidine mutant plasmids were prepared using the Q5 Site-Directed Mutagenesis Kit (E0552S NEB, Ipswich, MA). Gibson assembly (E2611S NEB, Ipswich, MA) cloning was used for both the loop chimera constructs and the pET22B PMO domain protein-expression constructs (for all alleles) according to the manufacturer’s instructions.

The generated plasmids were transformed into NEB 5-alpha Competent *E. coli* (NEB #C2987) for propagation and storage. Plasmid DNA was extracted with QIAprep^®^ Spin Miniprep Kit (Qiagen, Redwood City, California) (Cat. No. 27104) and linearized with *Nde*I (NEB, R0111S), *Ssp*I-HF^®^ (R3132S), or *Pci*I (NEB, R0655S) to transform *N. crassa*.

### Strain and culture conditions

The standard protocols for *N. crassa* growth, crosses and maintenance are available at the Fungal Genetics Stock Center website (http://www.fgsc.net/Neurospora/NeurosporaProtocolGuide.htm) (accessed 01/23/ 2019). All the strains used in this work are listed in Appendix 1-Table 4. The strains were grown in Vogel’s minimal medium (VMM)(Vogel, 1956) solidified with 1.5 % agar with supplements as required. For crosses, Westergaard’s synthetic cross-medium was used (Westergaard and Mitchell, 1947). The FGSC2489 laboratory strain was used as a parental strain and as a WT control for all experiments. The wild *N. crassa* strains used in this study were isolated from Louisiana, USA (Gonçalves et al., 2019; Heller et al., 2016; McCluskey et al., 2010; Palma-Guerrero et al., 2013).

### Transformation of *N. crassa*

Conidia of the auxotrophic histidine strain in the triple delete background (*ΔNCU01381ΔNCU01381NCU01382::hph; his-3^-^)* (Gonçalves et al., 2019) of *N. crassa* were transformed with linearized plasmid or PCR products by electroporation on a BIO-RAD Pulse controller plus and BIO-RAD gene pulser^®^ II. The conidia were electroporated using 1 mm gap cuvettes (BIO-RAD Gene Pulser ^®^/ MicroPulser™ Cuvette) (Bio-rad, Hercules, California) at 1.5 kV, 600 ohm, 25 μF as previously described (Margolin et al., 1997). For each transformation, around 30 (His^+^) prototroph transformants were selected and transferred to tubes containing VMM with Hygromycin B. The integration of the constructs in the selected transformants was confirmed by using F-130WH Phire Plant Direct PCR Kit (Thermo Fisher Scientific, Waltham, Massachusetts) and Sanger sequencing. Hygromycin B (10687010, 50mg/mL) (Thermo Fisher Scientific, Waltham, Massachusetts) was used at a final 200 µg/mL concentration for selection of transformants. Cyclosporin A (30024 Sigma, St. Louis, Missouri) was used at a final concentration of 5 µg/mL for selection. The L-histidine (L-Histidine hydrochloride monohydrate, 98%, Acros Organics, Waltham, Massachusetts) was used at a final concentration of 0.5 mg/mL to support the growth of his-auxotrophic strains.

### *N. crassa* crosses

Heterokaryotic transformants were used as a female parent strain and were grown on Westergaard’s synthetic cross-medium at 25 °C with constant light until protoperithecia were observed. The protoperithecia were fertilized with a conidial suspension from a male parental strain, FGSC9716/his^-^ or FGSC2489::GFP/his^-^ (FGSC2489*; csr-1::*P*ccg-1-gfp; his-3^-^*) and grown for an additional 10 days or until ascospores had been shot onto the lid of the Petri dish. To select homokaryons, the resulting ascospores were heat-shocked at 60 °C for 40 min and inoculated onto bottom agar plates (VMM with 1.5% agar and a mixture of 20% sorbose, 0.5% fructose, 0.5% glucose) and incubated at 30 °C overnight. The germinated ascospores were dissected using a stereomicroscope Stemi SV6 Zeiss and transferred to slants with VMM. The selection of the homokaryons was made depending on their (His^+^) prototrophy, Hygromycin B (Hyg^+^), and/or Cyclosporin A (Cyclosporin^+^) resistance. Additionally, PCR was done to confirm the presence of the construct using F-130WH Phire Plant Direct PCR Kit (Thermo Fisher Scientific, Waltham, Massachusetts) and Sanger sequencing.

### Germling-fusion assays

The strains were grown in VMM in slant tubes for four days at 30 °C; then, the tubes were placed at room temperature with constant light for 2 days. The conidia were resuspended in 2 mL of sterile ddH_2_O and filtered using cheesecloth. The styryl dye N-(3-triethylammoniumpropyl)-4-(6-(4-(diethylamino) phenyl) hexatrienyl) pyridinium dibromide (FM4-64, 514/670 nm absorption/emission Invitrogen™, Waltham, Massachusetts) was used at a final concentration of 16.5 μM in ddH_2_O from a 16.5 mM stock solution in DMSO, to stain germlings of *N. crassa.* An aliquot of 200 µL of the filtered conidia was stained with 40 µL of FM4-64 (16.5 µM) and incubated at room temperature for 15 min in the dark. The stained conidia were centrifuged in a microcentrifuge at 5000 rpm for 2 min, the supernatant was removed, and the conidia were washed twice with 1 mL of sterile ddH_2_O. The conidia were resuspended in 100 µL of sterile ddH_2_O and counted using the hemocytometer. The conidial suspension was adjusted to 3x10^7^ conidia/mL. An aliquot of 45 µL of the conidial suspension from the strain stained with FM4-64 was mixed with 45 µL of a conidial suspension from the strain that expresses cytoplasmic GFP. 80 µL of the final mixed suspension was spread on VMM agar plates (60 mm x 15 mm). Plates were incubated for 3.5 h at 30 °C. Agar rectangles of 3 cm x 2 cm approximately were excised and observed with a microscope ZEISS Axioskop 2 MOT. The microscope is equipped with an Illuminator Cool LED microscope, simply Better Control pE-300 white. The images were captured with a Q IMAGING FAST 1394 COOLED MONO 12 BIT microscope camera RETIGA 2000R SN:Q31594 (01-RET-2000R-F-M-12-C) viewed through a Ph3 40X/1.30 ∞/0.17 Plan-Neofluar oil immersion objective and then processed using iVision-Mac Scientific Image Processing Bio Vision Technologies (iVision 4.5.6r4). The fusion percentage of germlings was evaluated through observation of cytoplasmic mixing of GFP into germlings stained with FM4-64. Pictures for at least 15 fields were taken (for each filter DIC, Blue, GreenRed), depicting a total of at least 100 germling-contact events, with three biological replicates. Quantification is shown with error bars as SD. All the images were further processed using Fiji (ImageJ2, version 2.3.0/1.53f) (Schneider et al., 2012) Adobe Photoshop 2022 version 23.1.0, and Adobe Illustrator 2022 version 26.0.2. ANOVA statistical analyses were performed using GraphPad Prism 9. ANOVA statistical analysis was selected to assess differences in data between different strains/genotypes. One-way ANOVA for the analysis that involves one independent variable and two-way ANOVA for analysis with two independent variables was followed by Tukey’s test to perform multiple comparisons to assess significant differences. All p-values and CI 95% for each analysis are reported in P values source data.

### CWR-1 catalytic domain expression

Each *cwr-*1 allele was amplified from genomic DNA and cloned into the pet22B plasmid containing the pelB leader sequence. Each construct contained the PMO catalytic domain of the *cwr-1* gene with the signal peptide removed, followed by a C-terminal hexa-histidine tag with a thrombin cleavage site. The constructs were transformed into *E. coli* BL21 Ros2pLysS cells (QB3 Macrolab, Berkeley, California). Dense overnight cultures were grown in 100 µg/mL ampicillin (RPI), 20 µg/mL chloramphenicol (RPI) in LB and then 12 mL were added to 1 L of TB that contained 100 µg/mL ampicillin and 20 µg/mL chloramphenicol. At A600 of ∼0.6-0.8 IPTG was added to a final concentration of 100 µM, and cells were grown overnight at 18 °C. Each pellet was then resuspended in 1:4 weight/volume ratio of 50 mM Tris 30% Sucrose pH 8.0 1 mM AEBSF 1 mM benzamidine 0.1 mg/mL DNAase and 0.1 mg/mL lysozyme. This suspension was nutated for 30 min at 4 °C then centrifuged at 5000 x g for 55 min. The supernatant was decanted, and then the pellet was resuspended with 1:4 g/mL of ice-cold 5 mM Tris 5 mM MgCl_2_ pH 8.0. This suspension was nutated for 30 min at 4 ^°^C and then centrifuged at 8000 x g for 10 min. The sucrose and Mg supernatants were decanted and further clarified if needed by centrifuging at 4000 x g for 30 min and then combined. The combined supernatant was then run over a gravity column of His60 (Takara, Kusatsu, Shiga, Japan) resin. The column was washed with 20 column volumes (CV) of Buffer A (10 mM imidazole 50 mM NaPO_4_ 300 mM NaCl pH 8.0), then eluted with 5 CV of a stepwise gradient of 40 mM, 70 mM, 125 mM, and 250 mM imidazole. The buffer with 250 mM imidazole is herein referred to as Buffer B. The pure fractions determined by SDS-PAGE Stain-Free gel (BioRad, Hercules, California) were combined and concentrated to ∼1 mL and dialyzed using 7 KMWCO Snake Skin dialysis tubing (Thermo Scientific, Waltham, Massachusetts) into 20 mM Tris 150 mM NaCl pH 8.4. Thrombin agarose beads (15 µL) (Sigma, St. Louis, Missouri) were washed 3x with 500 µL of Buffer C (20 mM Tris 150 mM NaCl 2.5 mM CaCl_2_ pH 8.4), where the last wash of the resin was incubated for 30 min to activate the thrombin. The dialyzed protein was added to the beads and was gently rocked overnight at room temperature. The solution was centrifuged for 1 min at 1000 x g and then washed with 2x 1mL of Buffer B. The washes and supernatant were combined and run over a His60 bed. The resin was washed with 10 CV of Buffer A and 10 CV of Buffer B. The flowthrough and the Buffer A fractions were combined, concentrated, and dialyzed into 50 mM NaOAc 50 mM NaCl pH 5.5, then into 50 mM NaOAc 50 mM NaCl 5 µM CuSO_4_ pH 5.5, then into 50 mM MOPS 50 mM NaCl pH 7.0. The protein was then stored at 4 °C and showed negligible loss of chitin-degrading activity over several months.

### ICP-MS Cu quantification

Protein was diluted to 20 µM and digested in 2% nitric acid overnight at room temperature in 1.7 mL tubes (Starstedt, Nümbrecht, Germany). The samples were then centrifuged to remove insoluble material at 20,000 x g for 10 minutes, and the supernatant was transferred to new tubes. Samples were analyzed on a Thermo Fisher iCAP Qc ICP mass spectrometer in kinetic energy discrimination (KED) mode against a standard curve of known copper concentrations (CMS-5, Inorganic Ventures, Christiansburg, Virginia), with Ga (20 μg/L, Inorganic Ventures, Christiansburg, Virginia) as an internal standard. Each experiment was carried out in technical triplicate. Data are plotted with SEM error bars as calculated by GraphPad Prism 9.2.0.

### Purification of *N. crassa* cell walls

Cell wall material was purified as previously described(Maddi et al., 2009) with slight modifications: three 1 L flasks of 200 mL 1.5% agar 2% sucrose 1X Vogel’s salts were inoculated with conidia from FGSC2489, JW242, *Δcwr-1Δ81Δcwr-2,* and FGSC2489; *cwr-1*^D111^ ^HG5^ strains and allowed to grow at RT for 7 days with shaking (200 rpm). The conidia were separated from the mycelia with ∼200 mL of sterile MilliQ water and filtered through 4 layers of cheesecloth. The solution was then centrifuged at 1,200 x g for 5 min, washed by resuspending the conidia in 15 mL of PBS, and then centrifuged at 1,200 x g for 5 min. The conidia were then resuspended in PBS to a 1.0 × 108 conidia/mL titer. The conidia were lysed in a bead beater (BioSpec, Bartlesville, Oklahoma) with 0.1 mm glass beads for 5 x 1 min while resting on ice for 2 min in between cycles. Then the mixture was filtered using cheesecloth to remove the beads, and the disrupted conidia were resuspended in 15 mL of PBS. The solution was then centrifuged at 4,000 x g for 5 min, decanted, and resuspended in 15 mL of PBS with 10% SDS and centrifuged at 4,000 x g for 5 min. The supernatant was decanted, and conidia resuspended in fresh 15 mL of PBS with 10% SDS. The suspension was boiled for 15 min and cooled at RT. The solution was then centrifuged at 10,000 x g for 5 min, washed 2 times with 15 mL of PBS, and then 2 times with 15 mL MilliQ water. After gently pipetting away excess water, the disrupted conidia were frozen in liquid nitrogen and lyophilized overnight on a Labconco Freezone 2.5 plus Lyophilizer (Labconco, Kansas City, Missouri. The dried conidial cell wall material was brought to room temperature before use.

### Measurement of oxidized products

Chitin (80 mg) (α-chitin from powdered shrimp shells) (Sigma, St. Louis, Missouri) and β-chitin (Mahtani Chitosan Pvt. LTD. Veraval India) were dispersed and sonicated in 100 mM EDTA for 30 min and washed with MilliQ water by first resuspending with 1 mL of water and then centrifuging at 1000 x g for 10 mins. The supernatant was then discarded. The wash procedure was performed five times; then, the resulting chitin was resuspended to a stock concentration of 80 mg/mL. Substrates that did not show oxidative activity were tested at 20mg/mL included PASC that was prepared as previously described(Wood, 1988).Purified cell walls were resuspended in 50 mM MOPS pH 7.0. 10 µM PMO was incubated with 20 mg/mL of the substrate in 50 mM MOPS pH 7.0. The reactions were initiated with 1 mM ascorbate and incubated for 1 h, shaking at 37 °C. The samples were centrifuged at 20,000 x g at 4°C for 10 min, and the supernatant was transferred into new tubes to stop the reaction by removing from substrate. The products were analyzed with HPAEC-PAD using a Dionex ICS-5000 system (Thermo Fisher, Waltham, Massachusetts) with a gradient of three buffers: (A) 10 mM NaOH, (B) 100 mM NaOH, and (C) 100 mM NaOH 500 mM NaOAc. Buffers were prepared as dictated by the Thermo-Fisher manual, and the linear gradient (5 curve) was as follows 0.4 mL/min flowrate: 15%-50% B for 15 min; 50%-40% B 5%-60%C to 40% B 60% C for 20 min; isocratic 100% C for 5 min; then 15% B for 15 min. A gold electrode in carbohydrate-quadrupole mode was used as the detector. The autosampler was kept at 4 °C to preserve sample integrity. Each protein and substrate combination were analyzed at least in biological triplicate.

### Generation of C1-oxidized chitooligosaccharide standards

The standards were generated as previously described(Loose et al.,2014) with slight modifications. 3 mM of chitooligosaccharides (DP 3-6) (Megazyme, Wicklow, Ireland) were treated with 0.12 mg/mL of ChitO (Gecco Biotech, Groningen, The Netherlands) in 50 mM MOPS pH 7.0 overnight at room temperature and then aliquoted and stored at -20 °C until use.

### Liquid chromatography-tandem mass spectrometry

Samples of oligosaccharides were analyzed using a 1200 series liquid chromatography (LC) system (Agilent Technologies, Santa Clara, California) that was connected in line with an LTQ-Orbitrap-XL mass spectrometer equipped with an electrospray ionization (ESI) source (Thermo Fisher Scientific, Waltham, Massachusetts). The LC system was equipped with a reversed-phase analytical column (length: 150 mm, inner diameter: 1.0 mm, particle size: 5 µm, Viva C18, Restek, Bellefonte, Pennsylvania). Acetonitrile, formic acid (Optima LC-MS grade, 99.5+%, Fisher, Pittsburgh, Pennsylvania), and water purified to a resistivity of 18.2 MΩꞏcm (at 25 °C) using a Milli-Q Gradient ultrapure water purification system (Millipore, Billerica, Massachusetts) were used to prepare LC mobile phase solvents. Solvent A was 99.9% water/0.1% formic acid, and solvent B was 99.9% acetonitrile/0.1% formic acid (volume/volume). The elution program consisted of isocratic flow at 1% B for 5 min, a linear gradient to 95% B over 1 min, isocratic flow at 95% B for 4 min, a linear gradient to 1% B over 0.5 min, and isocratic flow at 1% B for 19.5 min, at a flow rate of 150 µL/min. The column compartment was maintained at 25°C, and the sample injection volume was 10 µL. Full-scan, high-resolution mass spectra were acquired in the positive ion mode over the range of mass-to-charge ratio (*m*/*z*) = 150 to 2000 using the Orbitrap mass analyzer, in profile format, with a mass resolution setting of 100,000 (at *m*/*z* = 400, measured at full width at half-maximum peak height, FWHM). For tandem mass spectrometry (MS/MS or MS^2^) analysis, selected precursor ions were fragmented using collision-induced dissociation (CID) under the following conditions: MS/MS spectra acquired using the Orbitrap mass analyzer, in centroid format, with a mass resolution setting of 15,000 (at *m*/*z* = 400, FWHM), normalized collision energy: 35%, activation time: 30 ms, and activation Q: 0.25. Mass spectrometry data acquisition and analysis were performed using Xcalibur software (version 2.0.7, Thermo Fisher Scientific).

### Electrospray Ionization Time of Flight Liquid Chromatography Mass Spectrometry (ESI-TOF LC-MS)

Acetonitrile (Optima grade, 99.9%, Fisher, Waltham, Massachusetts), formic acid (1 mL ampules, 99+%, Pierce, Rockford, Illinois), and water purified to a resistivity of 18.2 MΩꞏcm (at 25 °C) using a Milli-Q Gradient ultrapure water purification system (Millipore, Billerica, MA) were used to prepare mobile phase solvents S3 for LC-MS. Electrospray ionization mass spectrometry (ESI-MS) of proteins was performed using an Agilent 1260 series liquid chromatograph outfitted with an Agilent 6224 time-of-flight (TOF) LC-MS system (Santa Clara, CA). The LC was equipped with a Proswift RP-4H (monolithic phenyl, 1.0 mm × 50 mm, Dionex) analytical column. Solvent A was 99.9% water/0.1% formic acid (v/v) and solvent B was 99.9% acetonitrile/0.1% formic acid (v/v). PMOs were buffer exchanged into 25 mM ammonium bicarbonate buffer pH 7.5 using Biospin 6 (Bio-rad, Hercules, California) columns according to the manufacturer’s protocol and then spinning through an 0.22 µm cellulose acetate centrifugal spin filter. Samples of 1– 5 µL were injected onto the column, corresponding to 10 pmol of protein. Following sample injection, a 5–100% B elution gradient was run at a 0.30 mL/min flow rate over 8 min. Data were collected and analyzed by deconvolution of the entire elution profile (using Agilent Mass Hunter Qualitative Analysis B.05.00) to provide reconstructed mass spectra. Spectra were analyzed with open source Chartograph software (www.chartograph.com) (accessed 3/21/21) and compared to the mass predicted by Benchling (benchling.com accessed 3/21/21).

### O_2_ reduction measurements

The formation of H_2_O_2_ from PMO activity in the absence of polysaccharide substrate was measured using a horseradish peroxidase (HRP) coupled assay that oxidized Amplex™ red to produce a colorimetric readout. A PMO sample (2 µM) was incubated at room temperature with 100 µM Amplex™ red 1.3 µM HRP (Sigma), 2 mM ascorbate in 50 mM MOPS pH 7.0. The absorbance at 540 nm was measured at 30 s intervals for 10 min on SpectraMax340 spectrophotometer (Molecular Devices, San Jose, California). Each reaction was done in at least biological triplicate. Data are plotted with SEM error bars as calculated by GraphPad Prism 9.2.0.

### Homology modeling

The sequence of each haplogroup was used without the signal peptide as predicted by SignalP 5.0 (www.cbs.dtu.dk/services/SignalP/) (accessed 4/24/2021) and modeled in Swiss Prot using the copper-bound structure of *Ao*AA11 (PDB 4MAI) as the template. The resulting models were overlaid and visualized in ChimeraX 1.1.

### SSN generation and MSA analysis

The protein sequence of *cwr-1^HG1^* (NCU01380) was retrieved from the UniprotKB database and used as a search sequence in the EFI-EST website (Gerlt et al., 2015) (accessed 04/25/2021) using the blast option with a maximum of 10,000 hits and an e value of 5. These retrieved sequences (∼1700) were used to generate an SSN (sequence similarity network). An alignment score of 63 was used to generate and visualize the SSN in Cytoscape. The SSN data set is available in Appendix 1-Table 7. Source raw data for the SSN is available in the Figure 1-figure supplement 2 source data. ClustalW (accessed 04/25/2021) was used to generate a multiple sequence alignment (MSA) of the *cwr-1* haplogroup sequences and determine intra-haplogroup sequence ID%.

### EPR measurements

EPR spectra were acquired of 225 µM of CWR-1^HG1^ and 540 µM of CWR-1^H78A^ in 50 mM MOPS 10% glycerol pH 7.0 at 40 K. X-band continuous-wave (CW) EPR spectra were recorded on a Bruker ELEXSYS E500 spectrometer equipped with a cylindrical TE011-mode resonator (SHQE-W), an ESR 900-liquid helium cryostat, and an ITC-5 temperature controller (Oxford Instruments, Abingdon, United Kingdom). Spectra were acquired under slow-passage non-saturating conditions.

### Materials availability

All strains and plasmids listed in Appendix 1-Tables 1, 2 and 4 are available upon request or from the Fungal Genetics Stock Center (https://www.fgsc.net). Primers used in this study are listed in Appendix 1-Table 3. P value data for Figures 1, 3, 4, 5, 6 and Figure 1-figure supplement 3B are provided in the P value source data. Data for construction of the SSN is provided in Appendix 1, Table 7 and the raw data as Figure 1-figure supplement 2 source data.

**Figure 1.**
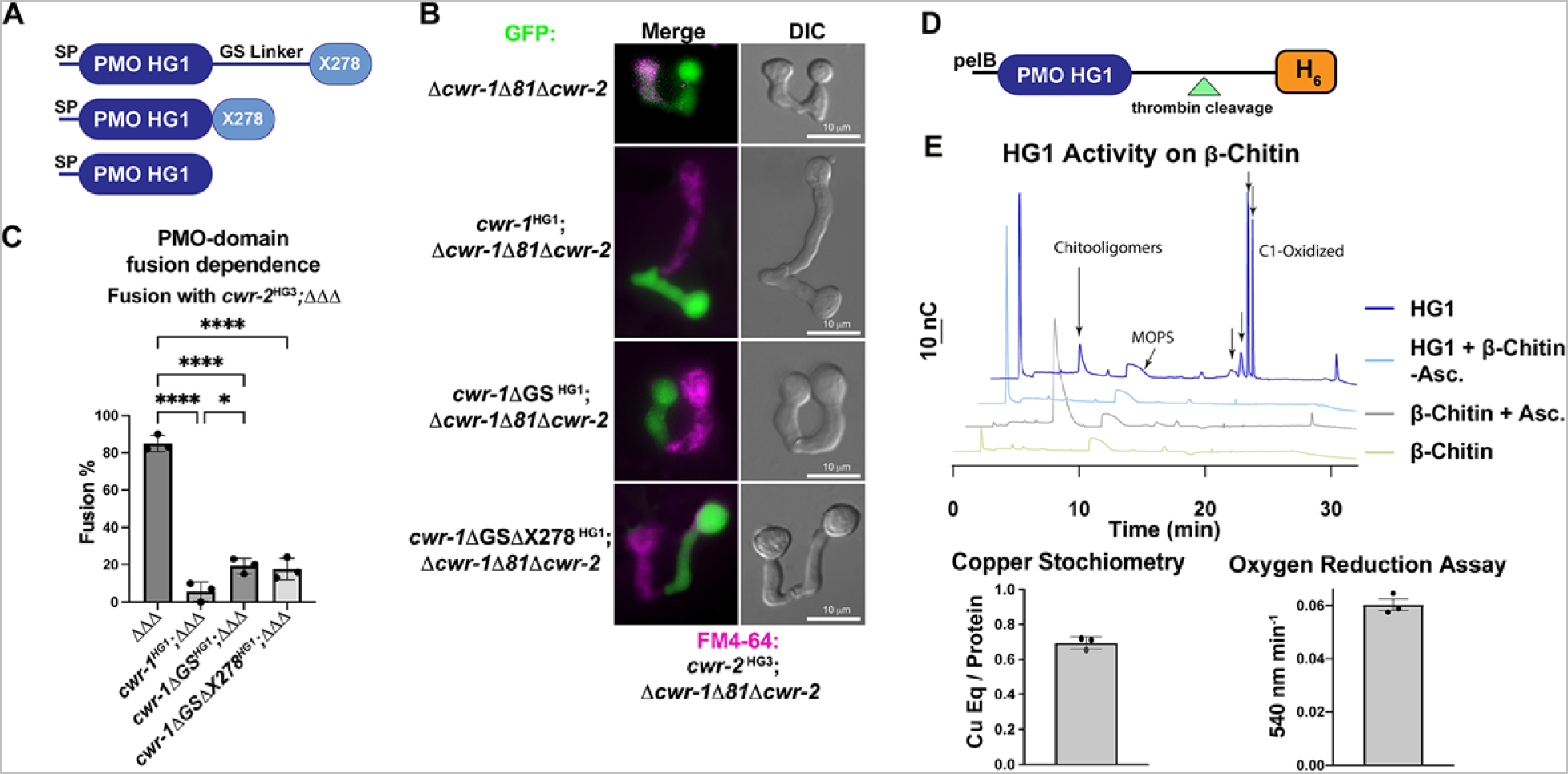
Characterization of the PMO domain from CWR-1 from a haplogroup 1 strain (FGSC2489) and functional dissection of CWR-1 domains *in vivo.* A) A schematic diagram depicting the series of truncated *cwr-1* constructs studied. SP indicates signal peptide; GS linker indicates the glycine/serine-rich region that connects the PMO catalytic domain to the presumptive chitin-binding module, X278. **B**) Cell fusion assays between *Δcwr-1Δ81Δcwr-2* germlings alone or bearing either HG1 *cwr-1* (FGSC2489) or truncated versions *cwr-1^Δ^*^GS^, *cwr-1^Δ^*^GS*Δ*X278^ (all expressing cytoplasmic GFP) and subsequently paired with FM4-64 stained *Δcwr-1Δ81Δcwr-2* germlings expressing a HG3 *cwr-2* allele (from JW258). (see Figure 1-figure supplement 3 for fusion rates with all HGs with HG1 and fusion controls) **C**) Quantification of cell fusion frequencies shown in Panel B of *Δcwr-1Δ81Δcwr-2* (GFP) germlings (*ΔΔΔ*), or *ΔΔΔ* germlings bearing HG1 *cwr-1^HG1^*or *ΔΔΔ* germlings bearing truncated versions *cwr-1^Δ^*^G/S^ or *cwr-1^Δ^*^GS*Δ*X278^ and paired with FM4-64 stained *ΔΔΔ* germlings expressing a HG3 *cwr-2* allele. Experiments were performed in biological triplicate, assessing fusion of 100 germling pairs for each replicate. A one-way ANOVA followed by Tukey post hoc test was used for statistical analysis, error bars represent SD, * p< 0.05, **** p<0.0001. Individual p-values are reported in P values source data. **D**) Schematic depiction of the *E. coli* expression constructs. pelB indicates the signal peptide, the PMO domain from a HG1 strain (FGSC2489), and a thrombin-cleavable hexahistidine tag showing cleavage site at the indicated triangle. (see Figure 1-figure supplement 4 for protein gel and MS of purified protein) **E**) Initial characterization of the PMO domain from HG1 strain (FGSC2489). This PMO exhibited C1-oxiding activity on chitin in the presence of ascorbate, reduced oxygen in the absence of chitin, and bound one copper atom per protein, all properties consistent with previously characterized AA11 PMOs. ICP analyses were done in technical triplicate, each datapoint in the oxygen reduction assay represents a biological replicate. All HPAEC-PAD assays were done in at least biological triplicate with a typical trace shown. Asc. is an abbreviation for ascorbate. Black arrows denote peaks that elute in the region of the method corresponding to C1-oxidized oligosaccharides. See Figure 1-figure supplement 5 for oxidized standards and Figure 1-figure supplement 6 for MS/MS spectra on PMO products from α-chitin.

**Figure 2.**
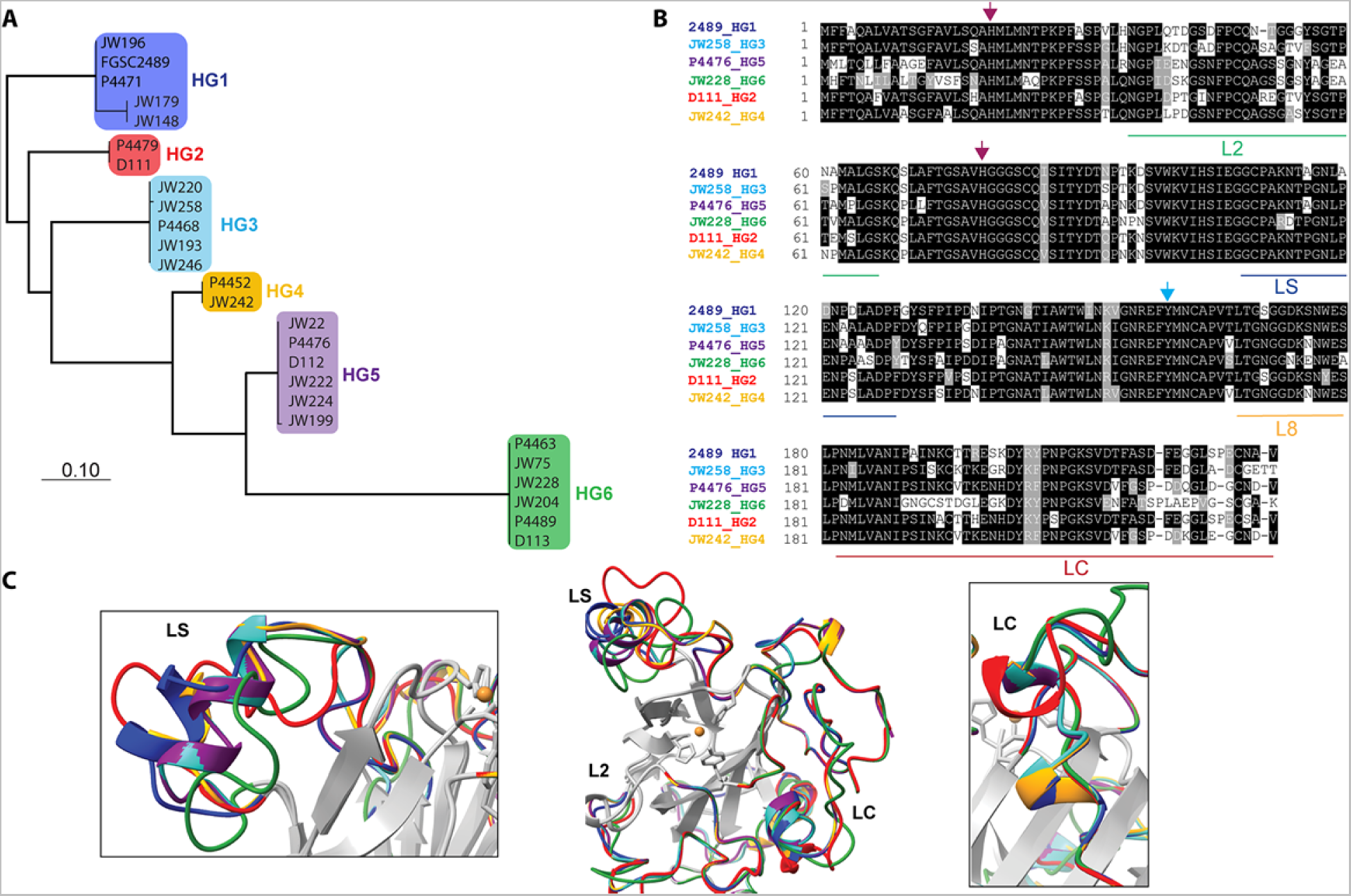
Comparison of CWR-1 PMO domains. In all panels, blue indicates *cwr-1* predicted proteins from HG1 isolates, red indicates HG2, light blue indicates HG3, orange indicates HG4, purple indicates HG5 and green indicates HG6 isolates. A). A phylogenetic tree was constructed using the predicted CWR-1 PMO domain from 26 wild-type isolates. The phylogenetic tree was made using PhyML Phylogenetic Maximum Likelihood and edited in MEGA11. B). Alignment of the six CWR-1 PMO domain protein sequences from representative isolates from HG1-6. The red arrows show the two histidine residues of the histidine brace that are involved in copper coordination. The blue arrow shows the position of the tyrosine involved in the secondary coordination sphere. L2, LS and LC correspond to loops that exhibited sequence variation between the CWR-1 PMO domains among the six different haplogroups. C). A SwissProt homology model of the PMO domain from the six haplogroups. There are apparent differences in the outer loops, whereas the core of the protein is not predicted to have significant differences between haplogroups. The four most affected loops are the loops that correspond to AA9 LS, L2, LC, and L8 loops (middle panel). A portion of the LS loop (left panel) has the most striking differences between haplogroups, and a portion of the LC loop (right panel) contains the second-best region for allelic differences.

**Figure 3.**
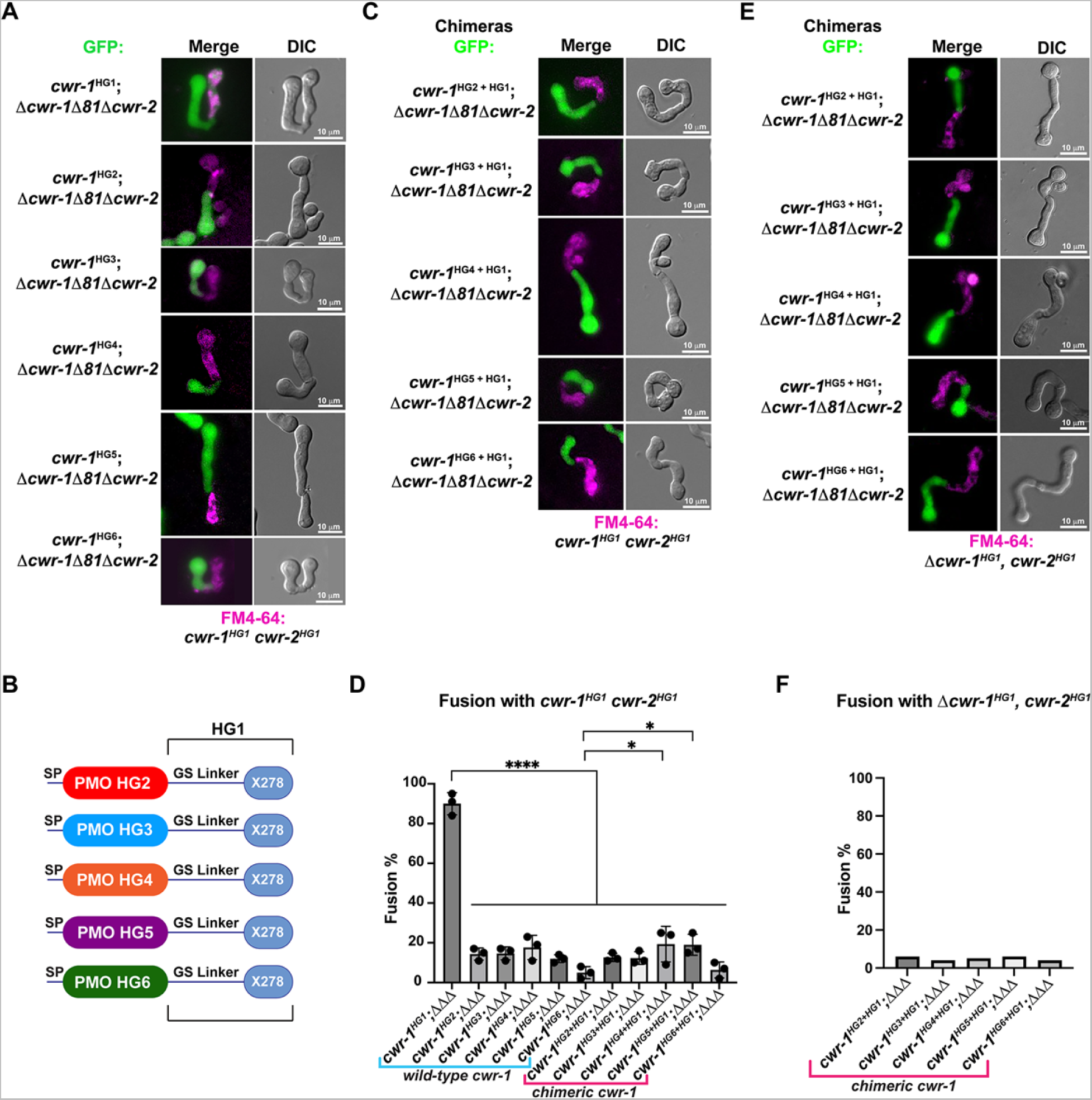
The PMO domain in CWR-1 functions to confer the allorecognition fusion checkpoint. A). Micrographs of the dominant fusion events between germlings expressing *cwr-1* alleles from each haplogroup in a Δ*cwr-1Δ81Δcwr-2* GFP background when paired with *cwr-1^HG1^ cwr-2^HG1^* germlings (FGSC2489) stained with FM4-64. B). CWR-1 chimeras with the PMO domain from the different *cwr* haplogroups (*cwr-1^HG2^* from D111, *cwr-1^HG3^* from JW258, *cwr-1^HG4^* from JW242, *cwr-1^HG5^* from P4476 and *cwr-1^HG6^* from JW228), with the glycine-serine linker and X278 domains from *cwr-1^HG1^* (from FGSC2489) are schematically shown. C). Micrographs of the dominant fusion events between germlings expressing the chimeric CWR-1 proteins paired with *cwr-1^HG1^ cwr-2^HG1^* germlings stained with FM4-64. D). Quantification of cell fusion frequencies depicted in panels A and C. The experiments were performed in biological triplicate, assessing fusion of 100 germling pairs for each replicate. A one-way ANOVA followed by Tukey post hoc test was used for statistical analysis, error bars represent SD, * p< 0.05, **** p<0.0001. Individual p-values are reported in P values source data. E). Micrographs of the dominant fusion event between GFP germlings expressing the chimeric CWR-1 proteins paired with a *Δcwr-1 cwr-2^HG1^* strain stained with FM4-64 F). Quantification of cell fusion frequencies between 100 germlings depicted in panel E. See Figure 3-figure supplement 1 for control experiments.

## Results

### The PMO domain of CWR-1 is sufficient to confer an allorecognition checkpoint

A multiple sequence alignment (MSA) of characterized AA11s revealed that three of the four characterized PMOs maintain a three-part architecture, while the other is a single domain PMO containing only a catalytic domain (Figure 1-figure supplement 1). The three domain architecture includes an N-terminus AA11 PMO domain (Hemsworth et al., 2014; Levasseur et al., 2013) followed by an extended GS-rich linker region that is likely disordered, and a C-terminal X278 domain proposed to be a chitin-binding domain; this domain is present in other predicted AA11 proteins and GH18 chitinases (Gonçalves et al., 2019; Hemsworth et al., 2014) (Figure 1A). AA11 proteins, similar to all other PMOs, contain a signal peptide at the N-terminus (predicted by SignalP) that directs the protein to the ER, and which is subsequently cleaved during translocation leaving a histidine residue at the N-terminus.

CWR-1 is a member of the three-region architecture cluster, and many fungi contain both AA11 PMO architectures in their genome. To more fully examine the relationships of predicted AA11 proteins, we performed sequence similarity network (SSN) analysis using the EFI-EST BLAST feature with the CWR-1^HG1^ sequence as the search peptide, which yielded ∼1700 sequences. The four characterized AA11 proteins that contained the GS linker and X278 domain were found in a single cluster (blue) (Figure 1-figure supplement 2). The single AA11 domain enzyme (*Af*AA11A) clustered separately into the red cluster. Since these domain architectures are clustered separately this suggests a different biological function in fungi. (Figure 1-figure supplement 2). This is not solely due to the difference in domain organization, for example starch PMOs with different architectures cluster together at ∼40% sequence ID (Vu et al., 2019). To determine which domain of CWR-1 was involved in allorecognition and the fusion checkpoint, a series of truncations were generated in the HG1 (FGSC2489) *cwr-1* allele: full-length *cwr-1*, removal of the GS linker, and removal of the GS linker and X278 domain (Figure 1A). Full length and truncated constructs were inserted at the *his-3* locus under the native promoter (1111 bp) and transformed into a strain of *N. crassa* bearing a deletion of *cwr-1* (NCU01380) and *cwr-2* (NCU01382) (*Δcwr-1Δ81Δcwr-2)* and expressing cytoplasmic GFP (Gonçalves et al., 2019). These strains were assessed for cell fusion in pairings with isogenic strains containing the *cwr-2* allele from each of the HG strains (HG1-6). The *cwr-1^HG1^* strain gave the strongest cell fusion block in pairings with *cwr-2^HG3^* germlings (Figure 1-figure supplement 3A, B). This *cwr-2^HG3^* strain was therefore used to test strains harboring the *cwr-1^HG1^* truncations that lacked the GS linker or the GS linker and the X278 domain. Strains containing any of the *cwr-1* truncations showed a block in cell fusion with the *cwr-2^HG3^*strain, including the truncation strain with only the catalytic PMO domain (Figure 1B, C). As a control, all of the truncation strains were also paired with a permissive mutant lacking *cwr-1* and *cwr-2* (*Δcwr-1Δ81Δcwr-2*); all showed robust cell fusion frequencies (Figure 1-figure supplement 3C, D). These data showed that the PMO catalytic domain was sufficient to cause cell fusion arrest at the CWR allorecognition checkpoint.

Given that the PMO catalytic domain alone was responsible for conferring allorecognition, we performed experiments to probe the activity for this domain. The PMO catalytic domain of the *cwr-1^HG1^* allele was expressed in *E. coli* using the the periplasmic expression pelB system which cleaves scarlessly during expression to ensure an N-terminal histidine with a C-terminal cleavable hexahistidine tag (Figure 1D) and subsequently purified using Ni-IMAC. The protein ran slightly higher (∼26-27 kDa) on an SDS-PAGE gel than the predicted size of 22.6 kDa (Figure 1-figure supplement 4A); the mass was validated with whole protein mass spectrometry. The spectrum revealed a deconvoluted mass of 22,624 Da, which corresponds to the exact mass of the protein with three disulfide bonds (22630 – 3 x (2 Da for each disulfide)) (Figure 1-figure supplement 4B). The CWR-1^HG1^ PMO domain bound ∼0.7 equivalents of copper after reconstitution and was shown to reduce oxygen to hydrogen peroxide in the presence of ascorbate (Figure 1E). The activity of the CWR-1^HG1^ protein was then tested with β-chitin as a substrate. New peaks between 18-22 min that corresponded to C1-oxidized products were only observed in the presence of both ascorbate and the CWR-1^HG1^ PMO domain (Figure 1E). C1-oxidized standards were generated from chitooligosaccharides (Figure 1-figure supplement 5A) using the chitin-C1-oxidizing enzyme ChitO (Figure 1-figure supplement 5B). The ChitO-C1-oxidized oligosaccharides eluted at timepoints similar to the new peaks from CWR-1^HG1^ PMO at around 18-22 min (Figure 1-figure supplement 5B, Figure 1E). To further validate the CWR-1^HG1^ C1 regioselectivity, tandem MS/MS was performed on the crude reaction from PMO^HG1^ action on α-chitin in the presence of ascorbate. The resulting spectra were consistent with C1-oxidation (Figure 1-figure supplement 6). These data were in line with all other characterized AA11 PMOs that exhibit C1-oxidative activity on both α-chitin and β-chitin.

Our data showed that the PMO domain of CWR-1^HG1^ was the driver for fusion arrest at the *cwr* allorecognition checkpoint. We therefore compared sequences of the PMO domain bioinformatically from the six different CWR-1 haplogroups. The phylogenetic tree showed identical clades to a tree created using full-length CWR-1 sequences (Gonçalves et al., 2019) (Figure 2A). The intra-haplogroup similarity was very high with >95% sequence identity across the PMO domain for alleles within a haplogroup (Appendix 1-Table 5). All six PMO haplogroups contained the residues essential for PMO activity; two histidines in the histidine brace and residues implicated in the catalytic mechanism, including a tyrosine involved in the secondary coordination sphere and other conserved hydrogen bonding residues (Span et al., 2017). Of note, the regions that have the most inter-haplogroup PMO domain differences corresponded to the AA9 nomenclature of LC, LS, and L2 loops, which have been shown to play a role in substrate binding and specificity. There were also minor differences in the L8 loop located opposite the substrate-binding surface of the PMO domain (Figure 2B). The differences between the different haplogroup PMO domains were more apparent when a homology model was constructed using the crystal structure of *Ao*AA11 (PDB: 4MAI) as a template (Figure 2C).

It is possible that the product of CWR-1 from each of the haplogroups was either generating a different product profile or recognized a different substrate. Mutations along substrate binding loops can change the regioselectivity of PMOs (C1 vs. C4) partially or entirely, and these residues can determine soluble substrate preferences (Courtade et al.,2018; Danneels et al., 2018; Liu et al., 2018; Vu et al., 2014). The two regions that exhibited the most prominent differences were the LC and LS loops, which was consistent with the hypothesis that a substrate/product(s) could be the signal for the *cwr* allorecognition checkpoint (Figure 2C).

Our *in vivo* data indicated that the PMO domain from *cwr-1^HG1^*was sufficient to confer the *cwr* allorecognition checkpoint. We therefore tested whether the full-length CWR-1 protein from each haplogroup was sufficient for fusion arrest and for conferring allelic specificity. Full-length *cwr-1* alleles from each haplogroup were expressed in a *Δcwr-1Δ81Δcwr-2* strain and confronted against *cwr-1^HG1^ cwr-2^HG1^*germlings. As expected, strains bearing *cwr-1* from any of the five different haplogroups showed significantly lower fusion rates as compared to a strain carrying *cwr-1*^HG1^ and paired with *cwr-1^HG1^ cwr-2^HG1^* germlings (Figure 3A, D). As a control to ensure fusion machinery is not impaired, strains bearing the six different *cwr-1* haplotypes showed high fusion rates with the *Δcwr-1Δ81Δcwr-2* strain (Figure 3-figure supplement 3A, C).

To determine whether the PMO domain from the CWR-1 proteins from all six different haplogroups was essential for allorecognition, chimeric constructs containing the N-terminal PMO domain from each HG (2-5) were fused to the GS linker region and X278 domain of *cwr-1^HG1^* (Figure 3B) in a *Δcwr-1Δ81Δcwr-2* strain (which also expressed cytoplasmic GFP). The resulting germlings were paired with a *cwr-1^HG1^ cwr-2^HG1^*strain, a Δ*cwr-1Δ81Δcwr-2* mutant, and a *Δcwr-1 cwr-2^HG1^*strain. The fusion percentages between germlings carrying any of the HG2-5 *cwr-1* chimeric constructs were low when paired with a *cwr-1^HG1^ cwr-2^HG1^* strain or with the *Δcwr-1 cwr-2^HG1^* strain, but were high when paired with the permissive *Δcwr-1Δ81Δcwr-2* mutant (Figure 3C, D, E, F; Figure 3-figure supplement 3B, C). No significant differences were observed in percentages of fusion when either full length *cwr-1* from the different haplogroups or with chimeric *cwr-1* alleles with the PMO domain from the different haplogroups were paired with a *cwr-1^HG1^ cwr-2^HG1^* germlings (Figure 3D). These experiments showed that allorecognition is mediated *in trans* between a cell containing *cwr-1* and a cell containing *cwr-2* from the different haplogroups and provide strong evidence that the PMO domain from the six different haplogroups conferred the allorecognition fusion checkpoint.

#### PMO catalytic activity is not required for the allorecognition fusion checkpoint

Our data above indicated that the PMO domain from each CWR-1 haplogroup was required to confer the allorecognition fusion checkpoint. These data suggested that the CWR-1 PMO domain from each haplogroup might generate a unique product distribution that would confer allorecognition. To test this hypothesis, the PMO domain from the remaining five CWR-1 haplogroups was expressed in a heterologous system, and the protein was purified using the same strategy for the expression of the PMO domain from the HG1 strain (FGSC2489) (Figure 1D, E). The CWR-1 proteins from HG2-6 ran slightly high (approximately 1-2 kDa higher than expected) on SDS-PAGE (Figure 1-figure supplement 4A) and were confirmed to be the correct mass (Figure 1 - figure supplement 4B), where each deconvoluted mass was consistent with three disulfides present on the protein (±1 Da) similar to the HG1 protein. All six PMOs from the different haplogroups oxidatively degraded both the α- and β-alloforms of chitin (Figure 4A, B), with β-chitin producing more oxidized fragments than α-chitin. Additionally, each of the six CWR-1 PMO proteins from the six different haplogroups generated C1-oxidized products, bound ∼1 equivalent of copper and reduced oxygen at similar rates (Figure 4C, D). None of the other substrates tested showed activity (Appendix 1-Table 6).

**Figure 4.**
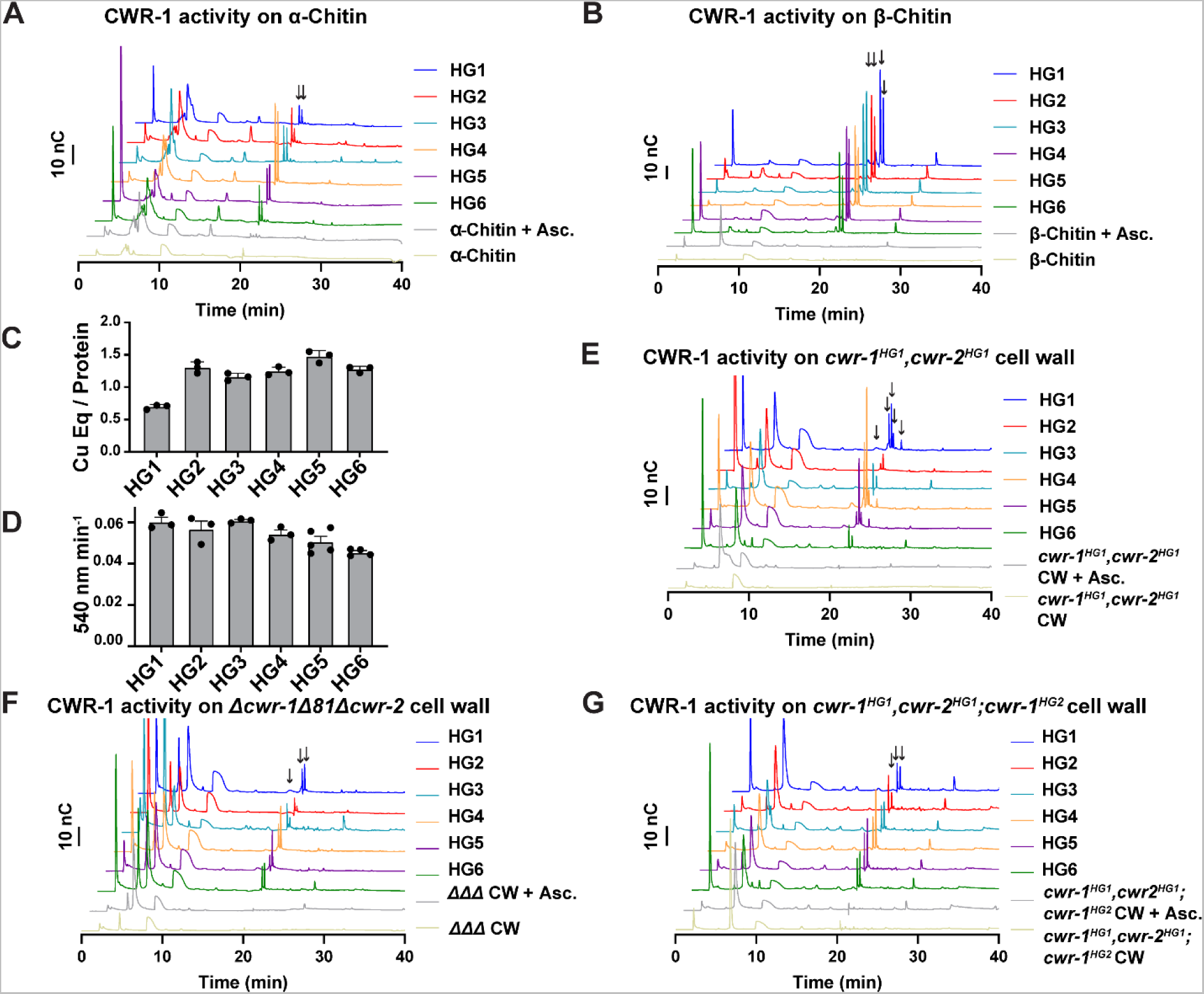
Catalytic activity of CWR-1 haplogroups. A-B) A comparison of the reaction products from CWR-1 from each of the six haplogroups on α-chitin and β-chitin. There were minor differences, but each haplogroup generates the same C1-oxidized fragments. C-D) Comparison of bound copper and oxygen reduction of CWR-1 from the six different haplogroups. All of the CWR-1 proteins bound ∼ 1 copper atom per PMO and reduced oxygen at similar rates. ICP experiments were done in technical triplicate, and oxygen reduction assays were done with each point representing a biological replicate. E-G) A comparison of reaction products of CWR-1 from each haplogroup on purified cell wall from the wild type HG1 strain (FGSC2489), the *Δcwr-1Δ81Δcwr-2* mutant strain, and a strain expressing a CWR-1^HG2^ PMO protein in a HG1 strain (*cwr-1^HG1^ cwr-2^HG1^*; *cwr-1^HG2^)* (Gonçalves et al., 2019). There were minor differences between the alleles and substrates, but all contain the same C1-oxidized chitin fragments. Asc. denotates ascorbate only. Black arrows denote peaks that elute in the region where C1-oxidized products elute. All HPAEC-PAD assays were done in at least biological triplicate.

Our *in vitro* data showed no selectivity differences between the CWR-1 PMOs from the six different haplogroups on α- and β-chitin alloforms. We therefore tested whether these HG1-6 PMOs had differences in activity on a more physiologically relevant substrate, the cell wall from a HG1 strain (FGSC2489). This cell wall was purified and assessed as a substrate for the PMO domains from the six different CWR-1 haplogroups. New peaks that corresponded to C1-oxidized chitooligosaccharides and soluble chitooligosaccharides appeared only in the presence of both reductant and the PMO. The products from the CWR-1 PMOs from haplogroups 1,4 and 5 showed a similar pattern of four peaks in the C1-oxidized chitin fragments region (Figure 4E), whereas the CWR-1 PMO from haplogroups 2,3 and 6 showed only two prominent C1-oxidized peaks (Figure 4E). Since this pattern did not correlate with phenotypic aspects of cell fusion incompatibility for the six CWR-1 haplogroups, we attributed this result to small rate differences between the CWR-1 PMO domains from the different haplogroups and heterogeneity in the purified cell wall suspension. As a control, cell wall components from the Δ*cwr-1Δ81Δcwr-2* strain were purified and used as a substrate. Although all six CWR-1 PMO haplogroups exhibited oxidized C1-oxidized products, there appeared to be two main groupings: CWR-1 from HG1 and 5 contained a third oxidized chitin fragment, while CWR-1 from HGs 2,3,4 and 6 showed only two prominent peaks (Figure 4F). These data indicated that cell walls from both HG1 and the *Δcwr-1Δ81Δcwr-2* strains were similar and that differences could be attributed to small rate differences or cell wall sample heterogeneity. Efforts to identify low abundance carbohydrates or other potential products through ESI-MS were unsuccessful.

PMO activity was also determined on purified cell walls from a self-incompatible strain (*cwr-1^HG1^cwr-2^HG1^*; *cwr-1^HG2^);* this strain shows reduced growth, asexual spore production and lack of self-fusion (Gonçalves et al., 2019). However, the pattern of products on this substrate for all six PMO domain haplogroups had very similar product profiles (Figure 4G). The PMO reaction products from CWR-1^HG1^ and CWR-1^HG4^ on chitin and purified cell walls from FGSC2489 (HG1) and JW242 (HG4) strains were also used in the germling fusion assays. However, neither the addition of the PMO reaction products nor the cell walls from the incompatible strains blocked or reduced cell fusion frequencies between compatible strains (Figure 4-figure supplement 1).

Since there were no striking differences between CWR-1 PMO haplogroups for either products or substrates, the requirement for catalytic activity in fusion blockage was tested. In *Serratia marcescens* and *Pseudomonas aeruginosa,* alanine variants in the second histidine (corresponding to His78 for CWR-1^HG1^) of the histidine brace in AA10 PMOs (CBP21 and CbpD, respectively), disrupt PMO activity with no oxidized products observed(Askarian et al., 2021; Vaaje-Kolstad et al., 2010). Therefore, we constructed *cwr-1^HG1^* and *cwr-1^HG6^* variants containing His à Ala substitution of the first histidine (His20), a second set of variants containing a His→Ala substitution at H78 and double His→Ala substitutions (H20A; H78A). Similar to the WT PMOs, the variant proteins’ deconvoluted mass was consistent with three disulfides and confirmed the construct was correctly expressed (Figure 5-figure supplement 1). These CWR-1 PMO His→Ala variants were assessed for catalytic activity, copper-binding, and/or subsequent hydrogen peroxide generation. All single and double histidine variants of PMO^HG1^, PMO^HG6^, and the TyràAla variant PMO^HG1^ were inactive on β-chitin, where no oxidative activity could be detected. (Figure 5A, B). PMO^HG1^ H20A and H78A variants and PMO^HG6^ H79A variants retained copper-binding after purification (Figure 5C). These variants were functional in an oxygen reduction assay but exhibited slightly lower rates as compared to WT CWR-1 PMO (Figure 4C). Both PMO^HG1^ and PMO^HG6^ His→Ala double variants greatly reduced copper-binding and had lower oxygen reduction rates (Figure 5D). X-band Continuous-wave EPR spectra were acquired for the WT PMO^HG1^ and PMO^H78A^ proteins. Both PMO^HG1^ and its H78A variants bound copper; this active site copper center is in the Cu(II) oxidation state under aerobic conditions. Cu(II) has an S=1/2 spin state, and EPR is a sensitive probe for the primary coordination sphere at the active site. The EPR spectra of the WT and H78A PMO^HG1^ variant proteins were compared to identify any differences in Cu(II) binding between WT and H78A mutant PMO proteins (Figure 5-figure supplement 2). The WT PMO^HG1^ spectrum is consistent with other reported PMOs (Frandsen et al., 2016) that contain axial Cu (II) sites with observed g values of g1 = 2.245, g2 =2.065, and g3 = 2.005 with visible N–hyperfine splitting. The spectrum from the PMO^HG1^ H78A variant is distinct from WT PMO^HG1^, displaying more splitting g2 and g3 values, indicative of a more rhombic coordination environment: g1 = 2.245 g2 = 2.08 and g3 = 2.05. The EPR differences between WT and the PMO H78A variant suggests a significant change to the primary coordination sphere of the Cu(II) ion and is presumably because the rigid histidine brace has been changed and replaced by water molecules. Since the PMO^HG6^ variant proteins were not substantially different from the PMO^HG1^ variant proteins, these results likely represent all the PMO proteins in the six haplogroups.

**Figure 5.**
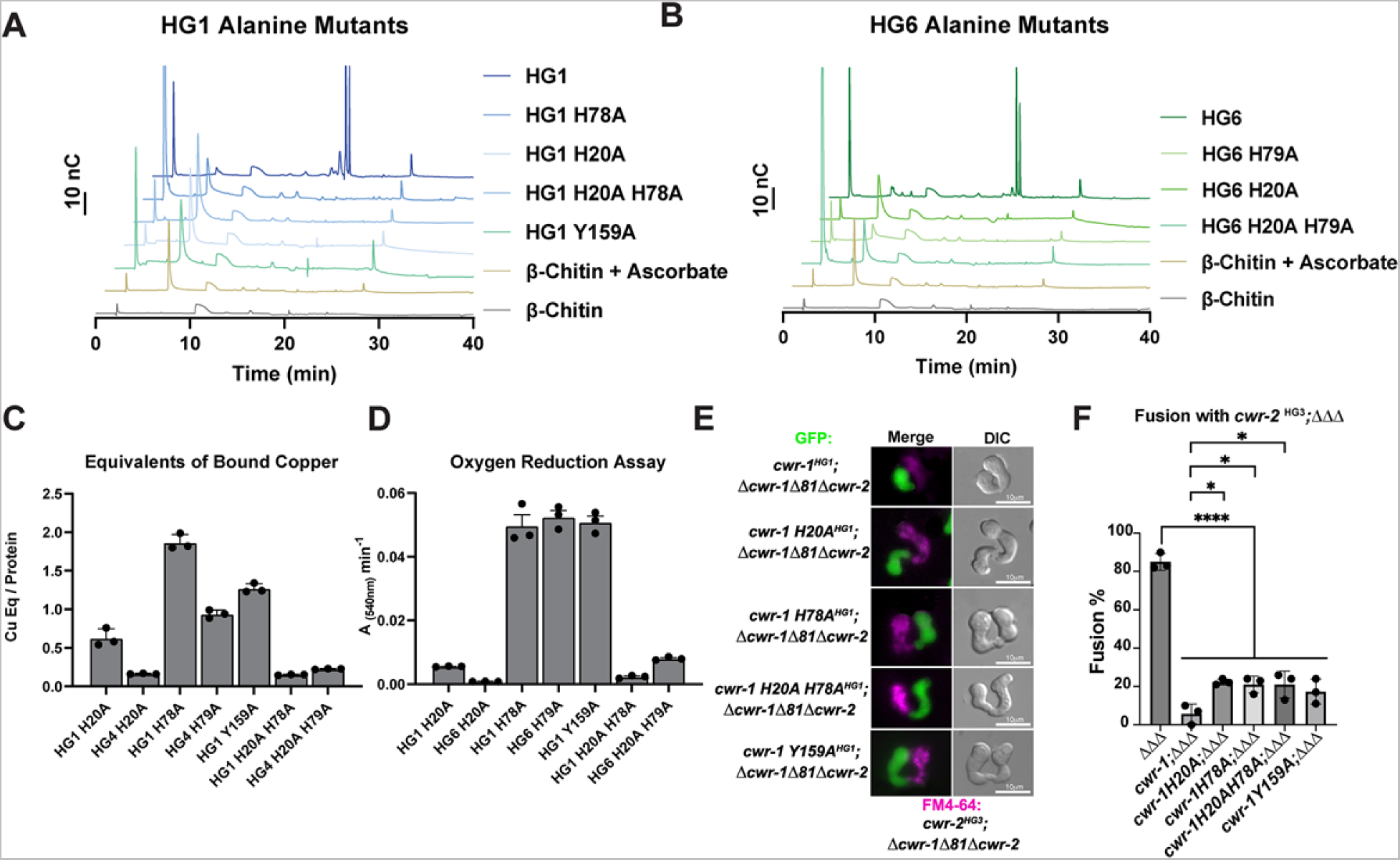
Characterization of CWR-1 PMO active-site mutants. A-B) Comparison of the ability of active-site variants of the PMO^HG1^ and PMO^HG6^ domains and their ability to degrade β-chitin. Mutation of one or both of the histidine brace residues abolished chitin-oxidizing activity. All HPAEC-PAD assays were done in at least biological triplicate. C) Comparison of bound copper of active site variants. Mutating the second histidine alone did not abolish copper-binding; however, mutation of the first histidine dramatically reduced the amount of copper bound, and mutation of both histidine residues almost eliminated copper-binding entirely. ICP experiments were performed in technical triplicate. D) Comparison of oxygen-reduction activity of active site mutants. All constructs that contained the first histidine to alanine variant had significantly reduced oxygen reduction activity. Each data point represents a biological replicate. E). Fusion test of the different CWR-1^HG1^ histidine variants and tyrosine variant paired with a strain expressing *cwr-2^HG3^* in the Δ*cwr-1Δ81Δcwr-2* mutant background. F). Quantification of cell fusion percentages depicted in panel E. The experiments were performed in biological triplicate, counting 100 germling pairs for each replicate. A one-way ANOVA followed by Tukey post hoc test was used for statistical analysis, error bars represent SD, * p< 0.05, **** p<0.0001. Individual p-values are reported in P values source data. The fusion percentages with *ΔΔΔ* and *cwr-1^HG1^; ΔΔΔ* are the same as in Figure 1C.

Previously, we tested whether a tyrosine residue in the PMO^HG1^ domain that is predicted to be important for catalysis was essential for allorecognition in pairings with a wild type strain (JW199, HG5); a statistically significant level of fusion was observed in these pairings, suggesting that Tyr159 was important for allorecognition (Gonçalves et al., 2019). Here, we further assessed these findings using an HG1 strain bearing a mutation in *cwr-1* (*cwr-1* ^HG1^ ^Y159A^) in a *Δcwr-1Δ81Δcwr-2* deletion strain and paired with isogenic germlings bearing a *cwr-2^HG3^*allele in a *Δcwr-1Δ81Δcwr-2* background. In these pairings, a significant block in cell fusion was observed (Figure 5E, F). These data suggest that other genetic factors in the JW199 strain affected cell fusion frequencies when paired with the HG1 *cwr-1*^Y159A^ mutant in an otherwise FGSC2489 background. However, using isogenic strains here showed that allorecognition and a significant cell fusion block occurred between a strain bearing only CWR-1^Y159A^ and a strain bearing only CWR-2^HG3^.

Mutations in the histidine brace resulted in chitin-inactive variants, although the purified PMO^HG1^ H20A, PMO^HG1^ H78A, and PMO^HG6^ H79A variants could still bind copper. With the retention of copper, hydrogen peroxide could still be generated from these single His→Ala variants. We, therefore, tested whether the single histidine brace variants (H20A and H78A) and the double His→Ala variant (H20A; H78A) were affected in allorecognition and cell fusion. Isogenic fungal strains were constructed bearing HG1 *cwr-1^H20A^*, *cwr-1*^H78A^, and *cwr-1* ^H20A; H78A^ alleles. Germlings from these strains were paired with germlings expressing *cwr-2*^HG3^ in the *cwr-1 cwr-2* deletion strain (*cwr-2*^HG3^; *Δcwr-1Δ81Δcwr-2*) that also expressed cytoplasmic GFP. Contrary to expectations, the single *cwr-1* histidine variants (H20A and H78A), including the inactive double mutant (H20A; H78A), still retained the ability to trigger allorecognition and block cell fusion when paired with germlings that expressed an incompatible *cwr-2^HG3^*allele (Figure 5E, F). As a control, all the His→Ala *cwr-1^HG1^* variants were paired with *cwr-1^HG1^ cwr-2^HG1^* (FGSC2489) germlings, and all underwent robust cell fusion (Figure 1-figure supplement 3C, D). These data rule out hydrogen peroxide generation as a contributing factor in blocking cell fusion, as the copper-deficient *cwr-1* ^H20A; H78A^ variant was not different from those that retained some copper-binding. Taken together, these data suggest that the *cwr* fusion block checkpoint is not dependent on enzymatic activity, and the CWR-1 PMO domain is not generating a signal to block cell fusion via a substrate preference or generated product.

### PMO domain loops confer specificity for cell fusion block

Our data showed that the PMO domain is essential for allorecognition, but that catalytic activity of the PMO domain was not required. An examination of a homology model of the six different CWR-1 PMO haplogroups based on the structure of *Ao*AA11 revealed that differences appeared to be most significant in the region of the LS loop (Figure 2C). To assess whether this region of CWR-1 was important for conferring allorecognition specificity, we constructed a loop-swap chimeric construct: the first where the entire LS loop and surrounding amino acids from PMO^HG1^ (I85 to G129) was replaced with the sequence from PMO^HG6^ (V86 to T130) (Figure 6A). If this region of the CWR-1 protein was necessary for allelic specificity, the chimeric strains would have a higher fusion frequency with a *cwr-2^HG6^* (in a *Δcwr-1Δ81Δcwr-2* background) strain and a lower fusion frequency with the *cwr-1^HG1^ cwr-2^HG1^*strain or *cwr-2^HG1^* in the *Δcwr-1Δ81Δcwr-2* background. However, the *cwr-1^HG1^LS ^HG6^* chimera maintained the same specificity as HG1 strain (Figure 6B, C). These studies indicate that this region, which is different in all six CWR-1 PMO haplogroups, was insufficient to switch allelic specificity. A second chimeric construct was generated; this new chimera includes LS loop, L8 loop, and the first half of the LC loop. The L8 and LC regions showed differences between haplogroups in the MSA and homology models, but more so in the case of the LC region. We hypothesized that a combination of these three regions could impart specificity. The region I85 to D201 of CWR-1^HG1^ was replaced with V86-D202 from CWR-1^HG6^ (Figure 6A). This *cwr-1^HG1/HG6^* chimera showed increased fusion percentages (∼55%) with *cwr-2^HG6^* cells, but a decreased fusion percentage (∼35%) with both the HG1 strain (FGSC2489) and a *cwr-2^HG1^* strain in the *Δcwr-1Δ81Δcwr-2* background (Figure 6D, 6E). These results suggest that the V86-D202 region in CWR-1 contains some but not all required residues to elicit the allorecognition response. To determine if we had created an allele with a new *cwr-1* specificity, we assessed fusion percentages of this chimera with strains bearing the other *cwr-2* haplogroup alleles in a *Δcwr-1Δ81Δcwr-2* background. The *cwr-1^HG1/HG6^*chimeric strain showed fusion percentages of ∼21%, ∼20% and ∼27% with strains bearing *cwr-2^HG2^*, *cwr-2^HG3^* and *cwr-2^HG5^*, but a higher fusion percentage with a *cwr-2^HG4^* strain (∼50%) (Figure 6E). Consistent with alterations in allelic specificity, these results were different from the *cwr-1*^HG1^ germlings paired with *cwr-2* germlings from the different haplogroups (Figure 6E). Thus, strains with the *cwr-1^HG1/HG6^* chimera showed the highest cell fusion percentages with the *cwr-2^HG4^* and *cwr-2^HG6^*strains but were not fully compatible with any of the strains containing *cwr-2* from the different haplogroups. These data and further studies into chimeric constructs of *cwr-1* in concert with chimeric constructs of *cwr-2* will help to define specific residues/regions required for haplogroup specificity and allorecognition that will enable the development of molecular models for somatic cell fusion arrest via the cell wall remodeling checkpoint.

**Figure 6.**
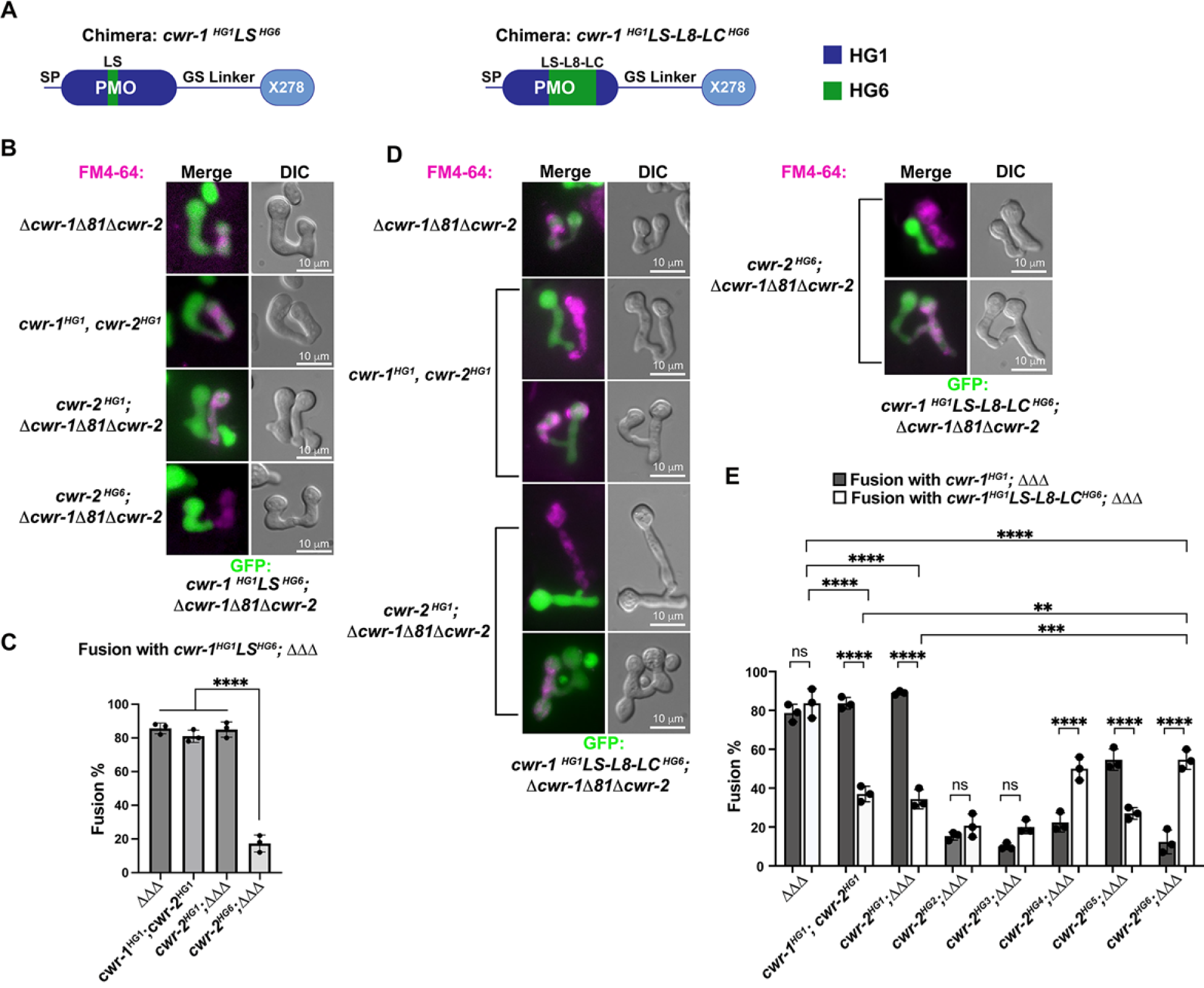
CWR-1 chimeras to define the PMO haplogroup specificity region. A) A schematic depicting where the loops from PMO^HG1^ domain that were replaced with loops from PMO^HG6^ (green). SP stands for signal peptide; GS linker stands for the glycine/serine-rich region that connects the catalytic domain to the X278, all of which were derived from CWR-1^HG1^. Navy is the region of the protein that is from PMO^HG1^, green is the region of the protein from PMO^HG6^. B) Cells expressing the chimera *cwr-1^HG1^LS^HG6^ ^(V86-T130)^* in a *Δcwr-1Δ81Δcwr-2* GFP strain were paired with indicated FM4-64 stained germlings. C) Cell fusion percentages between germlings depicted in panel B. The experiments were performed in biological triplicate, counting 100 germling pairs for each replicate. For statistical analysis a one-way ANOVA followed by Tukey post hoc test was used, error bars represent SD, **** p<0.0001. Individual p-values are reported in P values source data. D) Cells expressing the chimera *cwr-1^HG1^LS-L8-LC^HG6^ ^(V86-D202)^* in a *Δcwr-1Δ81Δcwr-2* GFP strain were paired with indicated FM4-64 stained germlings, with examples of blocked and fusing cells. E) Cell fusion percentages between the chimeric strain (*cwr-1^HG1^LS-L8-LC^HG6^*) and strains harboring the *cwr-2* alleles from all six haplogroups. Cell fusion percentages between *cwr-1*^HG1^ cells with strains carrying *cwr-2* alleles from different haplogroups were shown in Figure 1-figure supplement 3A, B. The experiments were performed in biological triplicate, counting 100 germling pairs for each replicate. For statistical analysis a two-way ANOVA followed by Tukey post hoc test was used, error bars represent SD, ** p< 0.01, *** p< 0.001 **** p<0.0001, ns: not significant.

## Discussion

The *N.* crassa cell wall remodeling checkpoint acts in *trans* via interactions between incompatible CWR-1 and CWR-2 during the process of cell fusion. If germlings or hyphae bearing *cwr-1* and *cwr-2* alleles of identical haplogroup specificity contact each other following chemotropic growth, cell wall dissolution is triggered, whereas cell fusion arrest occurs if strains carry *cwr-1* and *cwr-2* alleles of different haplogroup specificity. In this study, we analyzed cell fusion capability between engineered strains that were otherwise isogenic and harboring either *cwr-1* or *cwr-2* alleles from the distinct haplogroups or *cwr-1* chimeras. We determined that the PMO domain alone confers allelic specificity. However, our mutational, biochemical and *cwr-1* chimeric analyses showed that the PMO catalytic activity was not required for allorecognition and cell fusion arrest. These results suggest that CWR-1 and CWR-2 interact non-enzymatically in *trans* to trigger a block in cell wall dissolution. This interaction may include contact point(s) outside of the LS, LC, or L8 region of the CWR-1 PMO domain, as the chimeric constructs did not fully switch allelic specificity. This CWR-1 region may interact with the CWR-2 domains that face the cell wall or another, but unidentified, mediating partner may be important (Figure 7). Mechanistically, how CWR-1/CWR-2 interactions block in cell fusion is unclear, but we hypothesize that aspects associated with cell wall dissolution, including vesicle delivery or activation of cell wall remodeling enzymes could potentially be targets.

**Figure 7.**
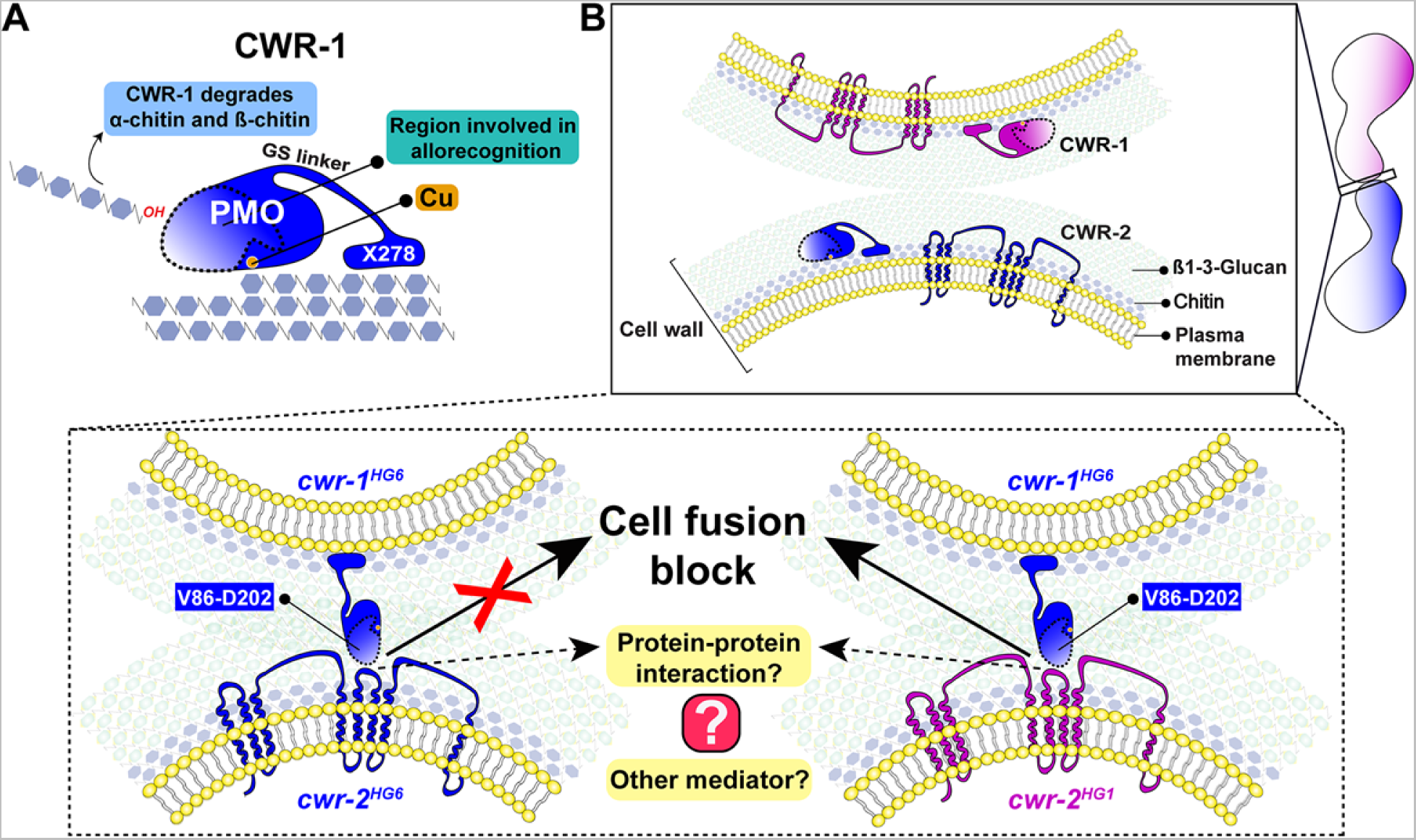
A schematic model that summarizes the role of CWR-1 and CWR-2 in allorecognition at the cell fusion checkpoint. A) The PMO domain, a glycine and serine-rich region and a putative chitin-binding module, X278 of CWR-1 are shown. The PMO domain degrades chitin and the region important for allorecognition is highlighted. B) The top panel shows a schematic showing the approach of the two tips of germlings before cell-cell contact. Once the germlings approach each other the bottom two panels show possible outcomes. In the left panel both germlings express a CWR-1 and CWR-2 protein belonging to the same haplogroup. This pair of germlings will not trigger the cell fusion block signal and will undergo cell fusion. In the right panel, the germlings express incompatible CWR-1 and CWR-2 proteins from different haplogroups therefore eliciting a cell-fusion blocking signal. The LC-LS-L8 (V86-D202) region of the protein is represented by the area inside the dotted line. It remains to be answered how CWR-1 and CWR-2 interact either directly through a protein-protein interaction or a mediator and which regions of CWR-2 are involved.

The CWR-1 PMO domain from all six haplogroups exhibits C1-oxidative activity on chitin and degrades β-chitin more efficiently than the α-chitin alloform, similar to the characterized, single-domain AA11 PMO (Støpamo et al., 2021). These data suggest that these two types of architectures in the context of chitin degradation may be redundant, which has been proposed previously for AA9 and AA13 PMOs (Vu et al., 2019). The biological role of the long GS linker and carbohydrate-binding module remains unclear. However, this linker and carbohydrate-binding module may allow the CWR-1 protein to remain bound to the cell wall through chitin-protein interactions, which may be important for non-allorecognition functions, including CWR-1 PMO activity. This work shows that CWR-1 can degrade chitin present in the fungal cell wall, which may aid in cell wall dissolution; the *Δcwr-1Δ81Δcwr-2* mutant shows a slight delay in fusion between compatible cells (Gonçalves et al., 2019) and is thus consistent with this hypothesis.

In biochemical analyses of the PMO domain catalytic function, it was surprising that several His→Ala variants still bound copper with 1:1 stoichiometry after reconstitution. Even though copper is bound, oxygen-dependent chitin turnover was abolished. The EPR of the bound copper showed the importance of the histidine brace in catalysis. Mutations of either the Cu-coordinating histidine or the non-coordinating axial tyrosine to alanine eliminated all catalytic activity on insoluble substrates. Previously, it was demonstrated that mutation of the axial tyrosine to phenylalanine in an AA10 PMO from *Thermobifida fusca* or an AA9 PMO from *Hypocrea jecorina* resulted in a less, but still active protein that maintains the ability to degrade phosphoric-acid swollen cellulose (PASC) (Kruer-Zerhusen et al., 2017; Jones et al., 2020). In nature, PMOs without tyrosine in this position often have phenylalanine as a replacement. It may be that any aromatic residue is essential in this position for oxygen-dependent catalysis and has some role in forcing the copper into the correct geometry through longer range structural interactions.

Peroxide-generating activity was observed in single CWR-1 histidine variants, yet subsequent Fenton-like chemistry that would cleave chitin did not occur. Thus, *in vivo,* the single histidine brace variants may retain the ability to bind copper and subsequently reduce oxygen, and while inactive on carbohydrate substrates, they will reduce O_2_ to H_2_O_2_ and may elicit an oxidative response. However, the block in cell fusion observed in germlings carrying single histidine brace CWR-1 variants suggests that H_2_O_2_ is not a signal involved in cell fusion. The increase in cell fusion percentages in strains carrying truncated versions of *cwr-1^HG1^*, or the histidine or tyrosine variants (∼20%) in pairings with *cwr-2* incompatible cells, as compared to fusion percentages in strains with wild type *cwr-1^HG1^*(∼6%) (Figure 1B, C 5E, F), may indicate that these changes may affect stability or alter CWR-1 conformational structure. Altogether, our data support that the catalytically dead CWR-1 variants still function to impose the cell wall remodeling checkpoint.

The region of the PMO domain essential for allelic specificity encompasses additional residues than what were swapped in the chimeric-loop constructs. As homology models may be inaccurate, changing alleles may require all three loops: L2, LS, and LC. More advanced structural studies would provide insight into the inter-haplogroup differences between CWR-1 proteins from the different haplogroups. The *cwr-1^HG1^ LS-L8-LC^HG6^* chimera showed a reduction in cell fusion percentages when paired with strains expressing *cwr-2* from any of the six different haplogroups (Figure 6E). These data suggest that the CWR-1^HG1/HG6^ chimera expresses a functional protein capable of allorecognition and that this chimera may confer a novel *cwr-1* specificity. This *cwr-1^HG1^ LS-L8-LC^HG6^*chimera will be useful in defining domains of CWR-2 that confer haplogroup allelic specificity, and raises the possibility of constructing a new *cwr* haplogroup by the construction of a *cwr-2* chimera that is fully compatible with strains carrying the *cwr-1^HG1^ LS-L8-LC^HG6^* chimera. It should be noted, however, that wild isolates contain both *cwr-1* and *cwr-2* alleles, and thus potentially two incompatible *cwr-1-cwr-2* interactions can be triggered during the cell fusion process.

There are several examples of proteins that have been co-opted by biology to perform a completely different role in the cell. This behavior is termed “moonlighting” (Jeffery, 1999, 2018) and includes enzymes that form structural components of mammalian eyeball lenses (Piatigorsky, 1998), an aconitase that acts as an iron sensor (Gunawardena et al., 2016), and at least one other PMO *e.g.* the mucin binding protein/ AA10 PMO, GbpA, from *Vibrio cholerae*, which is important in binding to the host cell (Bhowmick et al., 2008; Loose et al., 2014). There has also been a recently characterized chitin-degrading PMO from *Pseudomonas aeruginosa,* CopD, that is involved in immune evasion. However, the CopD PMO active site appears to be essential for this effect as histidine to alanine variants disrupt the immune evasion phenotype (Askarian et al., 2021). PMOs are known to have another protein-protein interaction partner, cellobiose dehydrogenases (CDHs), that serves as an electron source for catalysis (Kittl et al.,2012; Laurent et al., 2019; Tan et al., 2015). Bioinformatic searches yielded no portion of CWR-2 that contains a consensus sequence for flavin, heme, or metal sites, but this does not rule out entirely that the CWR-2 proteins are not electron donors. However, this observation could explain why all of the PMO domains from the six different HGs have the same substrate and products, as each reaction cannot occur until reducing equivalents are delivered. Since the only domain present in CWR-2 is an uncharacterized domain of unknown function, future experiments should be done to rule out this possibility.

Since three of the four characterized AA11s are so similar in both sequence and architecture (Figure 1-figure supplement 1, 2), it may be the case that these three characterized AA11s are also functioning in the same biological role as CWR-1. RNA-seq experiments show *cwr-1* transcripts are upregulated during starvation (Wu et al., 2020) and fruiting body development(Zheng et al., 2014), which may suggest that *cwr-1* has two biological roles. Allorecognition loci in filamentous fungi are under balancing selection, with defined haplogroups in population samples (Gonçalves et al., 2020). Since many fungal species do not have multiple genomes sequenced and available in online databases, and many others do not have facile genetic methods, it is unclear how widespread the role of *cwr-1/cwr-2* is in allorecognition. However, there are several other species that have linked *cwr-1/cwr-2* loci that fall into haplogroups in population samples, including *Fusarium verticillioides, Fusarium fujikuroi*, *Trichoderma harzianum, Zymoseptoria tritici,* and *Fusarium oxysporum* (Gonçalves et al., 2019). Future work will define regions of allelic specificity in CWR-1 and corresponding regions of CWR-2 and probe mechanistic aspects of how the interactions/functions of these proteins mediate the cell wall remodeling checkpoint.

## Acknowledgments

We want to thank the Glass laboratory and Marletta laboratory members for critical reading of the manuscript. We acknowledge Daniel Brauer for help in analyzing the whole protein mass spectra. We kindly thank the C. Chang laboratory for the use of their ICP-MS. We want to acknowledge the QB3 mass spectrometry facility and Anthony T. Iavarone for his help in analyzing high-resolution mass spectra of the reaction products. The QB3/Chemistry Mass Spectrometry Facility received support from the National Institutes of Health (grant 1S10OD020062-01). A.M.R.-R., N.L.G., and partially T.C.D. were supported by a National Science Foundation grant (MCB-1818283). T.C.D. was also partially supported by the National Science Foundation grant CHE-1904540.

## Author contributions

T.C.D., A.M.R.-R., M.A.M., and N.L.G. designed the study and experiments. T.C.D. and A.M.R.-R. prepared the original manuscript. T.C.D., A.M.R.-R., R.I.S., A.P.G., M.A.M, and N.L.G. edited and revised the manuscript. T.C.D. expressed and purified protein, purified *N. crassa* cell wall, performed ICP analysis, generated oxidized standards, performed HPAEC-PAD analysis, oxygen reduction experiments, analyzed mass spectra, and performed bioinformatic analyses. T.C.D. and A.M.R.-R. performed the homology modeling analysis. A.M.R.-R. and T.C.D. constructed plasmids. A.M.R.-R. and A.P.G. generated *N. crassa* strains. A.M.R.-R. performed germling-fusion microscopy analysis. R.I.S. performed EPR measurement and analysis. M.A.M. and N.L.G. provided supervision and project administration.

---

**Figure 1-figure supplement 1.**
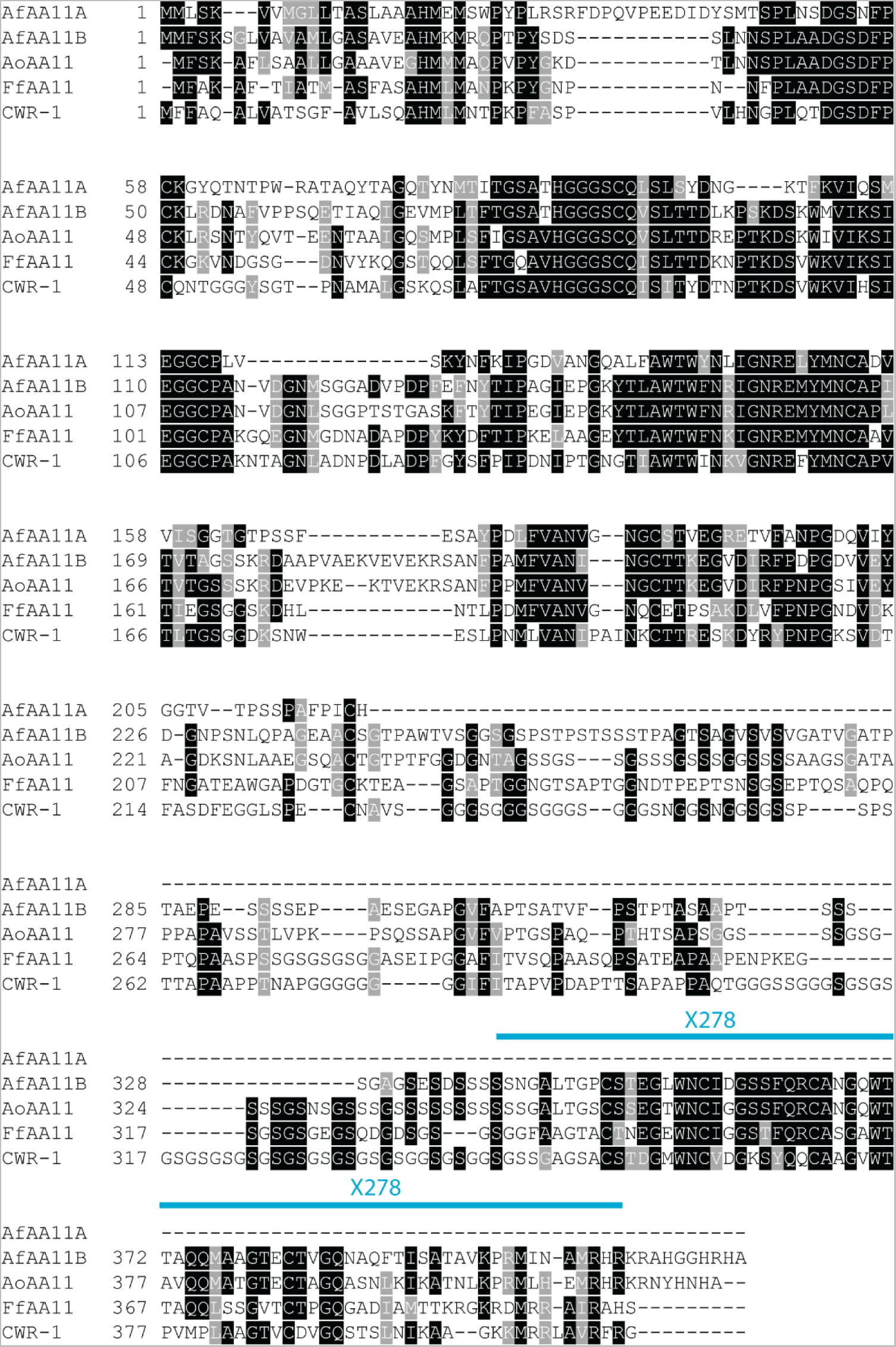
Alignment of characterized AA11s. All of the AA11s besides AfAA11A have a very similar architecture to CWR-1, including the GS linker and X278 domain.

**Figure 1-figure supplement 2.**
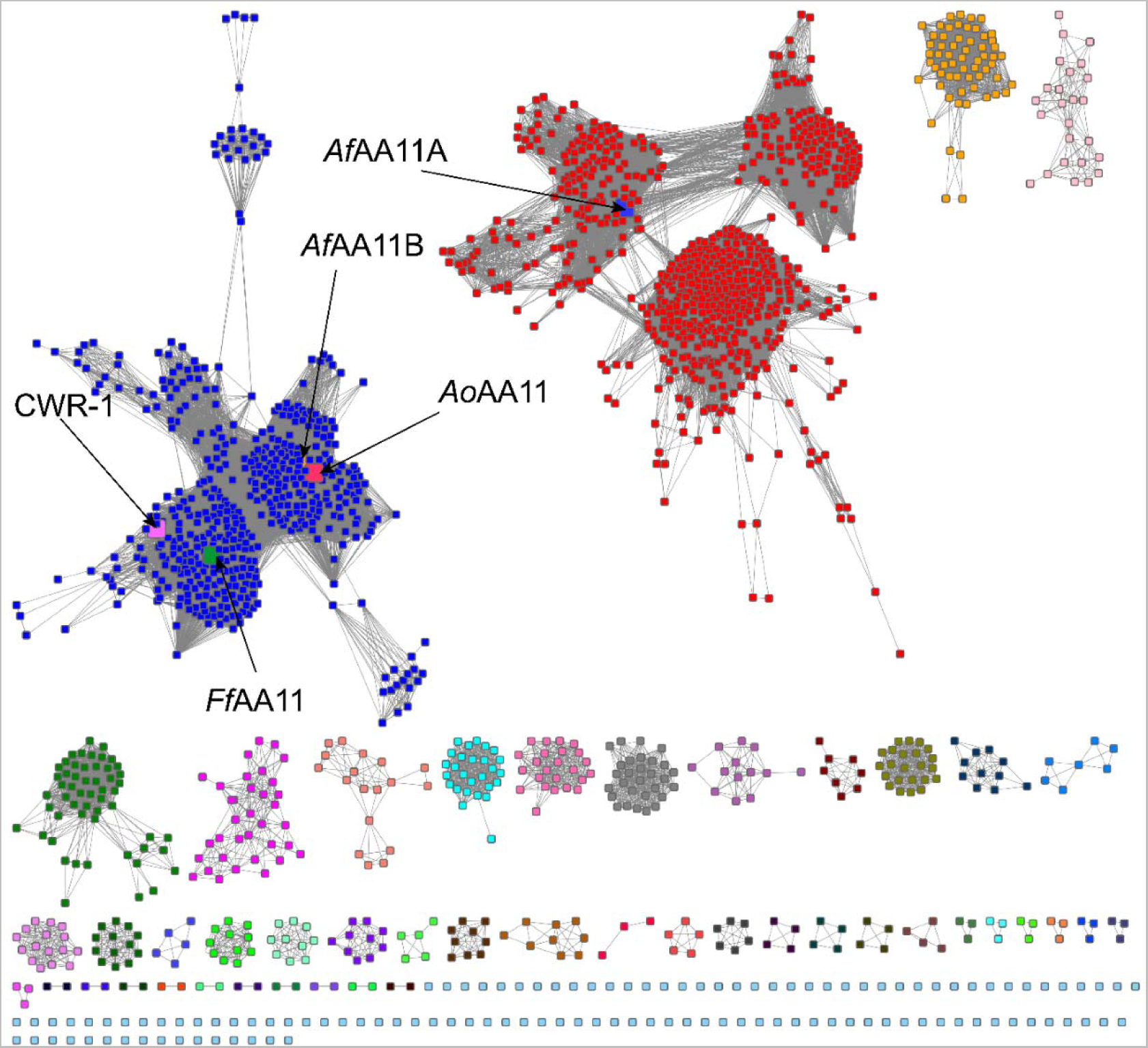
AA11 sequence similarity network (SSN) annotated with characterized proteins. This SSN was generated using the EFI-EST BLAST feature with the CWR-1^HG1^ sequence as the search peptide. This search yielded approximately 1700 sequences (Appendix 1, Table 7) which were visualized using an alignment score of 63 in this network corresponding to ∼40% sequence ID. The five labeled nodes are the characterized AA11s. The four found in the blue cluster contain the GS linker and X278 domain, while *Af*AA11A is a single domain enzyme consisting of only the AA11 domain. The CWR-1 and *Ff*AA11 PMOs cluster away from the others at a higher alignment score and may have a unique role within the blue cluster. The other clusters are proteins with similar sequences and presumably similar functions, but do not contain characterized proteins characterized. Some species contain genes that belong to several clusters, but most often the red and blue clusters. Raw data used for SSN construction is provided in Figure 1-figure supplement 2 source data.

**Figure 1-figure supplement 3.**
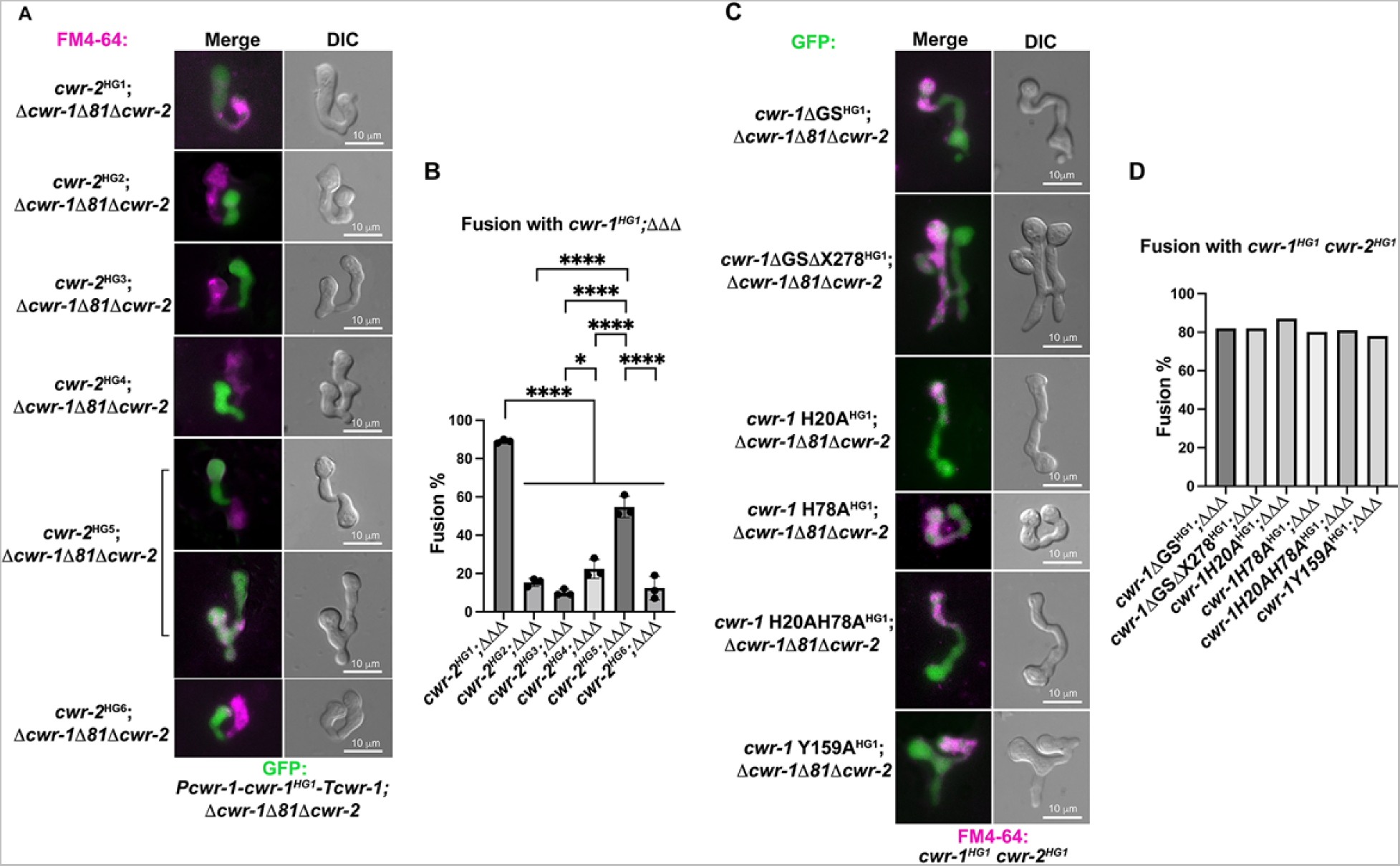
Germling fusion phenotype and percentages among engineered stains. A) Pairings between a *cwr-1*^HG1^ strain and strains carrying *cwr-2* alleles from the six different haplogroups. A strain harboring a *cwr-1*^HG1^ allele and expressing cytoplasmic GFP (*his-3::*P*cwr-1-cwr-1^HG1^-*T*cwr-1*; *Δcwr-1Δ81Δcwr-2::hph; csr-1-Pccg-1-gfp*) was paired with strains carrying *cwr-2* alleles from the different haplogroups (HG2-HG6) (*his-3::* P*tef-1-cwr-2^HG-X^*-V5-T*ccg-1*; *Δcwr-1Δ81Δcwr-2*) stained with FM4-64; fusion percentages were determined microscopically. B) Quantification of cell fusion percentages depicted in panel A. The experiments were performed in biological triplicate, assessing fusion outcome of 100 germling pairs for each replicate. For statistical analysis a one-way ANOVA followed by Tukey post hoc test was used, error bars represent SD, * p< 0.05, **** p<0.0001. Individual p-values are reported in P values source data. C) Strains carrying *cwr-1^HG1^*alleles bearing a deletion of the glycine-serine rich (GS) domain (*cwr-1^ΔGS^*) or the GS linker and the X278 domains (*cwr-1^ΔGSΔX278^*), the catalytic domain mutants (*cwr-1^H20A^, cwr-1^H78A^, cwr-1^H20A;H78A^* and *cwr-1^Y159A^*, all in a *Δcwr-1Δ81Δcwr-2* background and expressing cytoplasmic GFP, were paired with isogenic *cwr-1^HG1^cwr-2^HG1^*germlings stained with FM4-64. The genetic background for all the *cwr-1* mutants is *his-3::*P*cwr-1-cwr-1^MUTANT^*-Tcwr-1; *Δcwr-1Δ81Δcwr-2::hph; csr-1-Pccg-1-gfp*. D) Quantification of cell fusion percentages between 100 germling pairs depicted in panel C.

**Figure 1-figure supplement 4.**
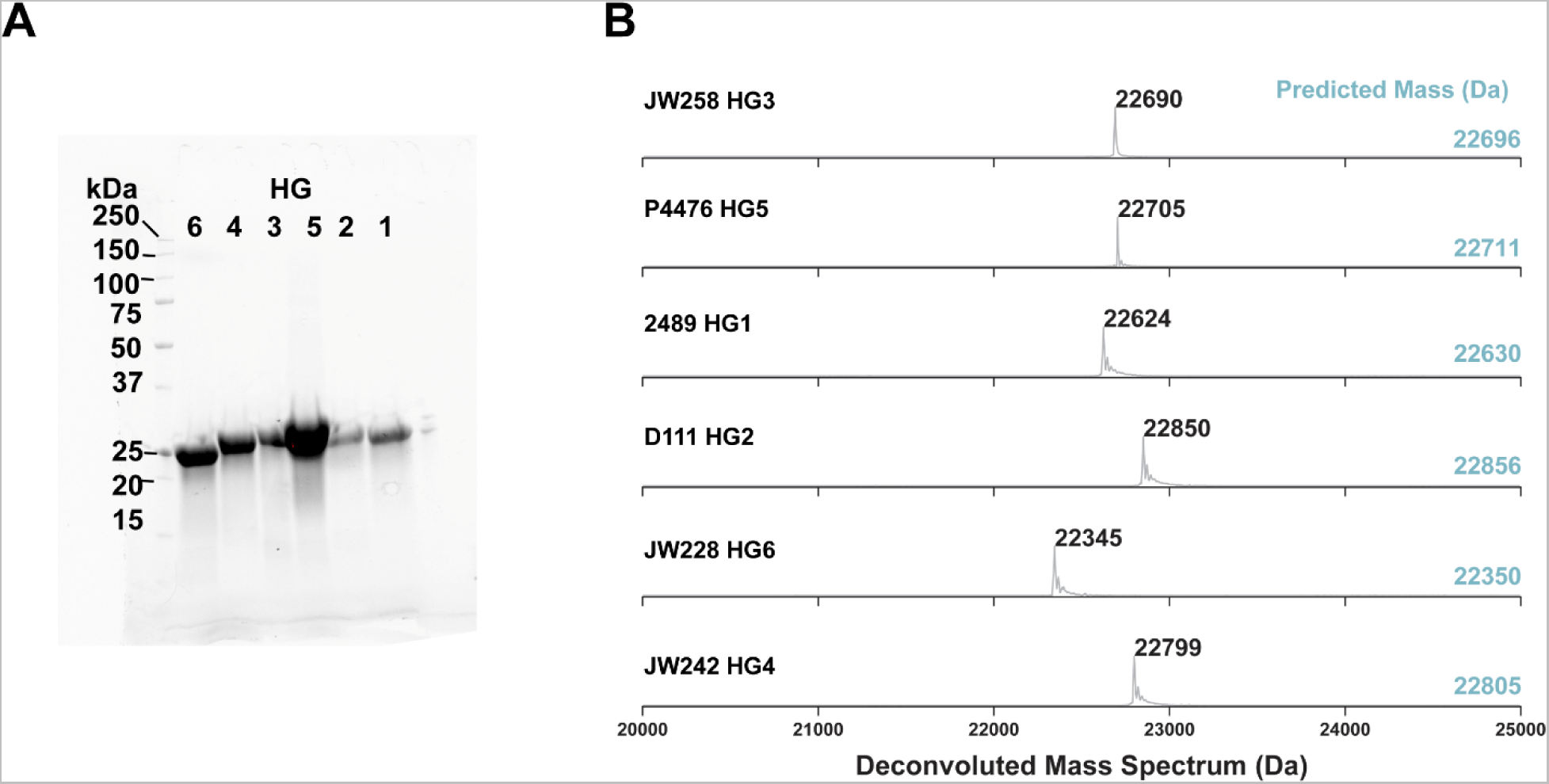
Purification and size of PMO domains of CWR-1 haplogroups (HG) proteins. A) An SDS-PAGE gel showing purity of CWR-1 PMO domains encoded by *cwr-1* alleles from the six different haplogroups. The haplogroup is denoted above each lane. Original data provided in Figure 1-figure supplement 4A source data. B). Deconvoluted protein mass spectrum of purified wild type PMOs between 20 kDa and 25 kDa. The predicted mass of each PMO is shown on the right in teal. Each mass observed is 6 or 5 Da lower than predicted, consistent with three pairs of disulfide bonds present in these PMOs. The haplogroup (HG) of each PMO domain and the strain it was derived from are noted on the left. The HG for A and B correspond

**Figure 1-figure supplement 5.**
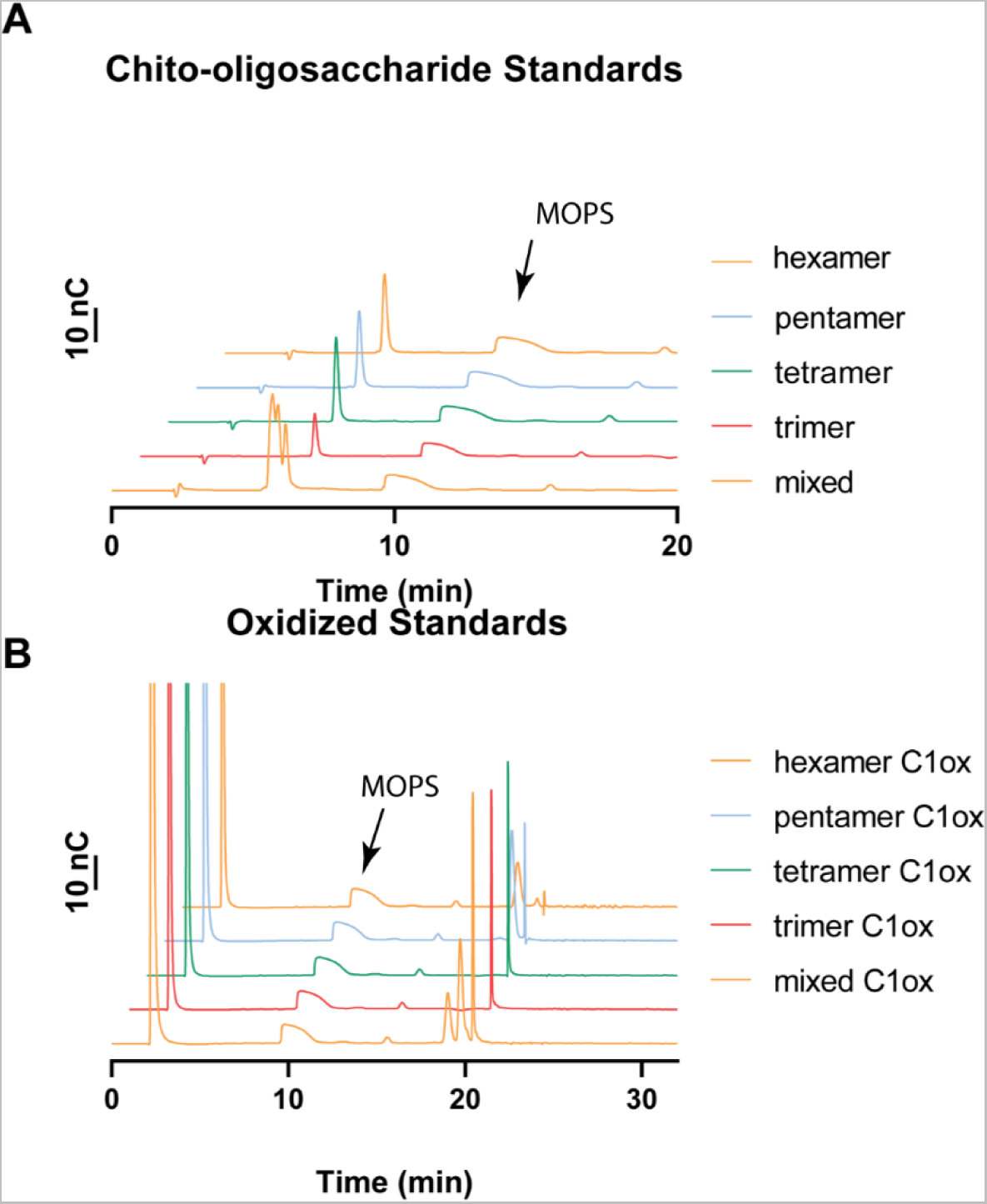
Oligosaccharide standard chromatograms and elution times. A) Chito-oligosaccharide elution times. Chito-oligosaccharides elute early on in the HPAEC-PAD trace. The tetramer and trimer elute at very similar times. An arrow denotes where the buffer, MOPS, elutes as a tailing peak. B) C1-oxidized chitooligosaccharides (ChitO treated chitooligosaccharides) elute ∼18-22 min. The trimer and tetramer elute at very similar retention times. There is degradation in the form of smaller oxidized oligosaccharides present in the pentamer and hexamer samples. An arrow denotes where the buffer elutes.

**Figure 1-figure supplement 6.**
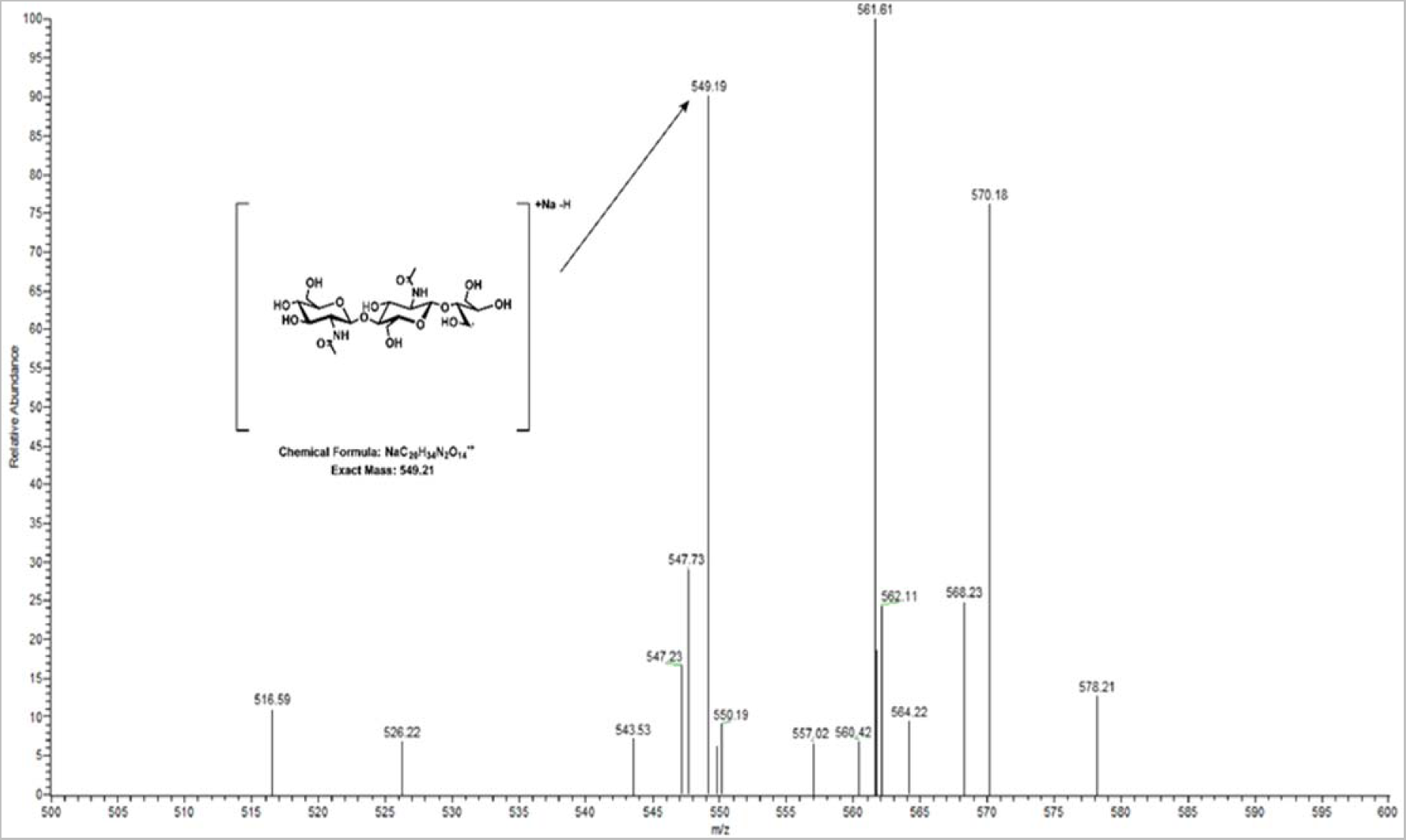
MS/MS fragment is consistent with C1 oxidation. A MS/MS spectrum from the product of PMO^H**G1**^ treated α-chitin. This protein was used as a representative sample for the others as the products do not appear different by HPAEC-PAD between the PMOs from the six different haplogroups. The inset shows an expanded region of the fragmented oxidized trimer ion with an exact mass of 666.2327. This precursor ion was chosen as it was the most abundant oxidized oligosaccharide ion in the MS run. There is some ambiguity between C1 and C4 fragments, however one peak can be assigned as a fragment consistent with C1 and not C4 oxidation.

**Figure 3-figure supplement 3.**
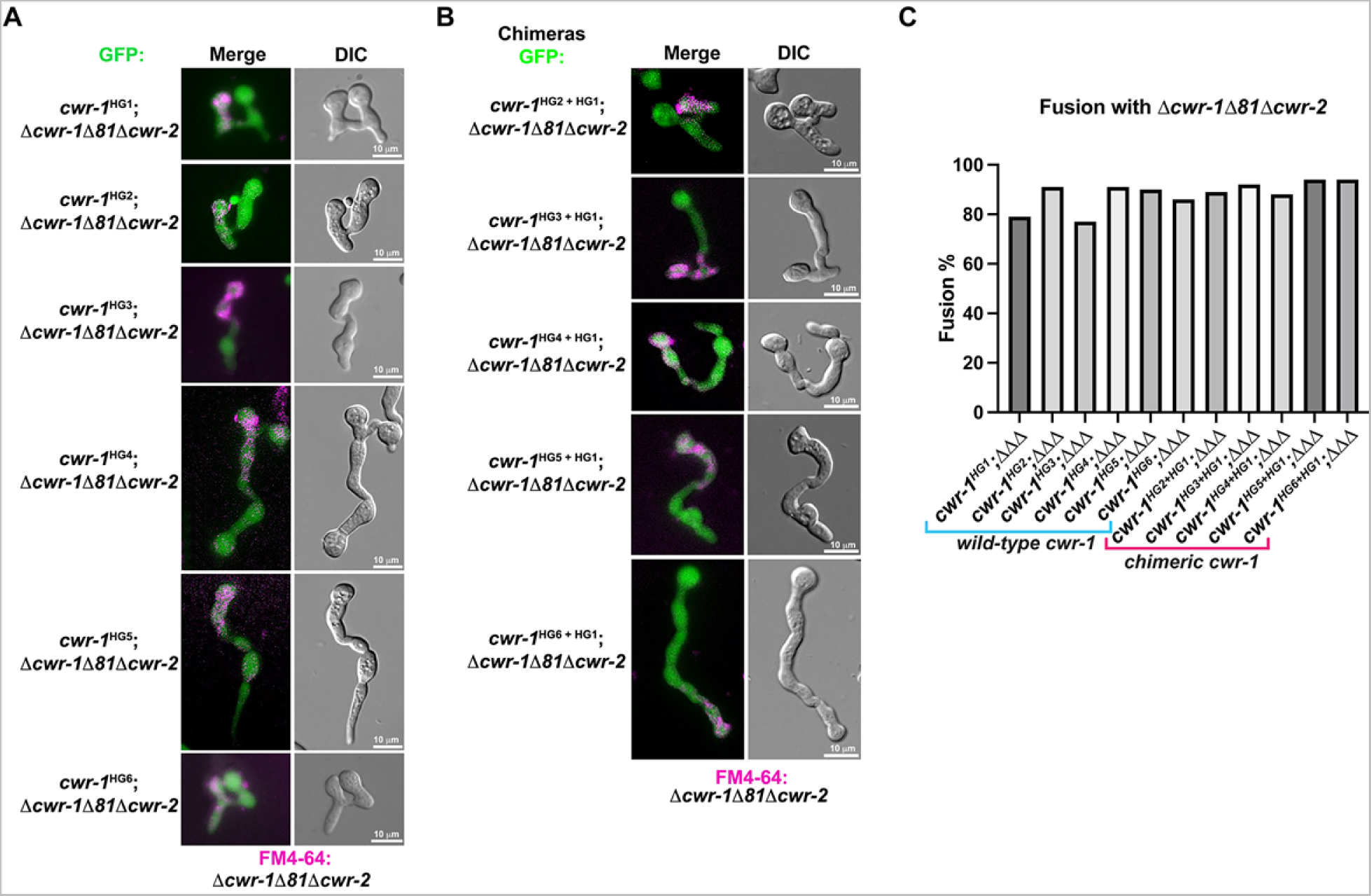
Germling fusion phenotype and fusion percentages in strains bearing *cwr-1* from the different haplogroups and *cwr-1* chimeras. A). Representative micrographs of fusion phenotypes of cells expressing *cwr-1* alleles (from HG1-6) and paired the permissive *Δcwr-1Δ81Δcwr-2* mutant stained with FM4-64. **B)**. Representative micrographs of *cwr-1* chimeras containing the PMO domain from *cwr-1* alleles from HG1-6 in *Δcwr-1Δ81Δcwr-2* GFP background and paired with a *Δcwr-1Δ81Δcwr-2* mutant stained with FM4-64. **C).** Quantification of cell fusion percentages between 100 germling pairs depicted in panels A and B.

**Figure 4-figure supplement 1.**
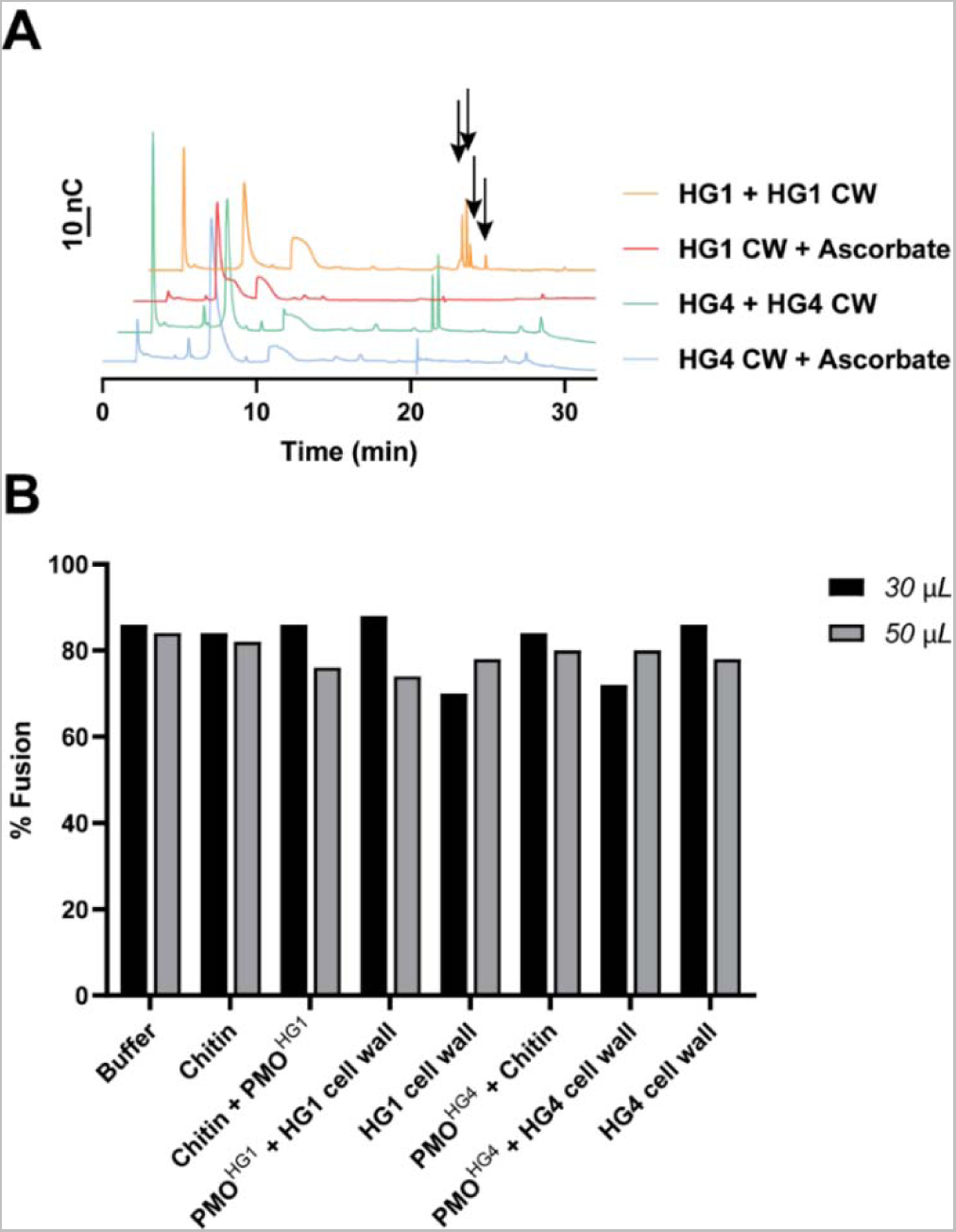
HPAEC-PAD traces of reaction before fusion experiment and cell fusion percentages. A) Purified CWR-1 PMO (from HG1 or HG4) was mixed with conidial cell wall sample, with or without ascorbate. All solutions used reagents at the same concentration as other activity assays (1 mM ascorbate, 50 mM MOPS pH 7.0, 10 µL PMO, 20 mg/mL substrate) and purified PMO^HG1^ or PMO^HG4^. The traces with ascorbate present show that chitin degradation has occurred in both cases with new peaks that appear between 18-20 min in the sample with PMO present consistent with PMO degradation of chitin from the cell wall. Black arrows denote peaks that elute in the 18-20 min window where C1 oxidized products elute. B) Crude reaction mixtures from the enzymatic activity of the HG1 or HG4 purified PMO acting on either chitin or purified conidial cell walls from HG1 or HG4 strains was obtained; the reaction was allowed to proceed for 1h, insoluble material was removed through centrifugation at 10,000 x g for 10 minutes. The resulting supernatant was used in a germling fusion assay between *Δcwr-1* (stained with FM4-64) and *Δcwr-1* germlings expressing cytoplasmic GFP to determine if addition of reaction products affected fusion percentages. Quantification of cell fusion frequencies between 50 germling pairs and represented in percentages. 30 and 50 µL denote the amount of completed reaction mixture added.

**Figure 5-figure supplement 1.**
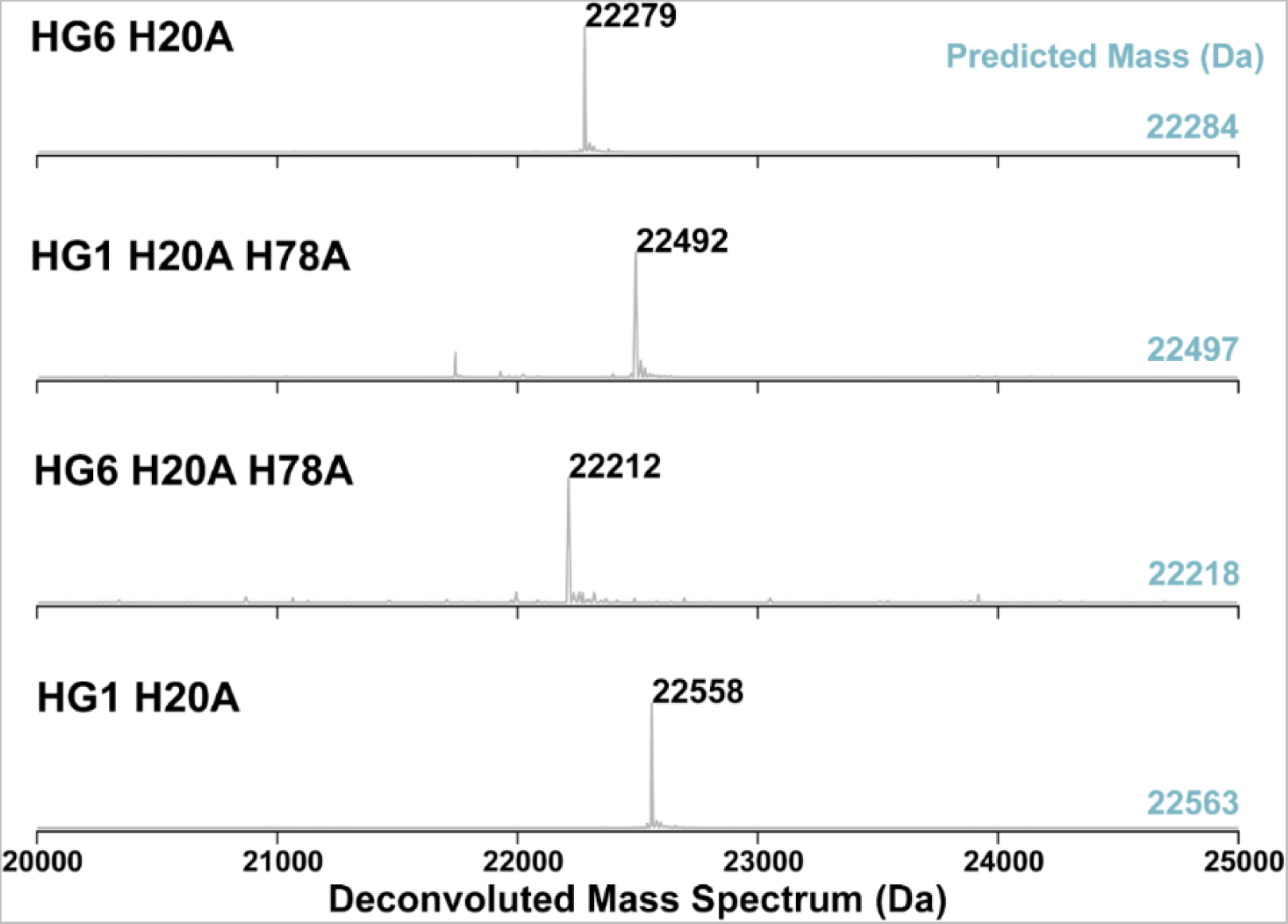
Deconvoluted mass spectrum of PMO histidine mutants. These PMO^HG1^ variants have a histidine to alanine substitution of the first histidine (His20; the first reside after the signal peptide), a second variant containing a His◊Ala substitution at H78 and double His◊Ala substitutions (H20A; H78A). All proteins were processed correctly in the pelB system. The predicted mass of each PMO^HG1^ variant is shown on the right in teal. Each mass observed is 6 or 5 Da lower, consistent with three pairs of disulfide bonds present in these PMOs.

**Figure 5-figure supplement 2.**
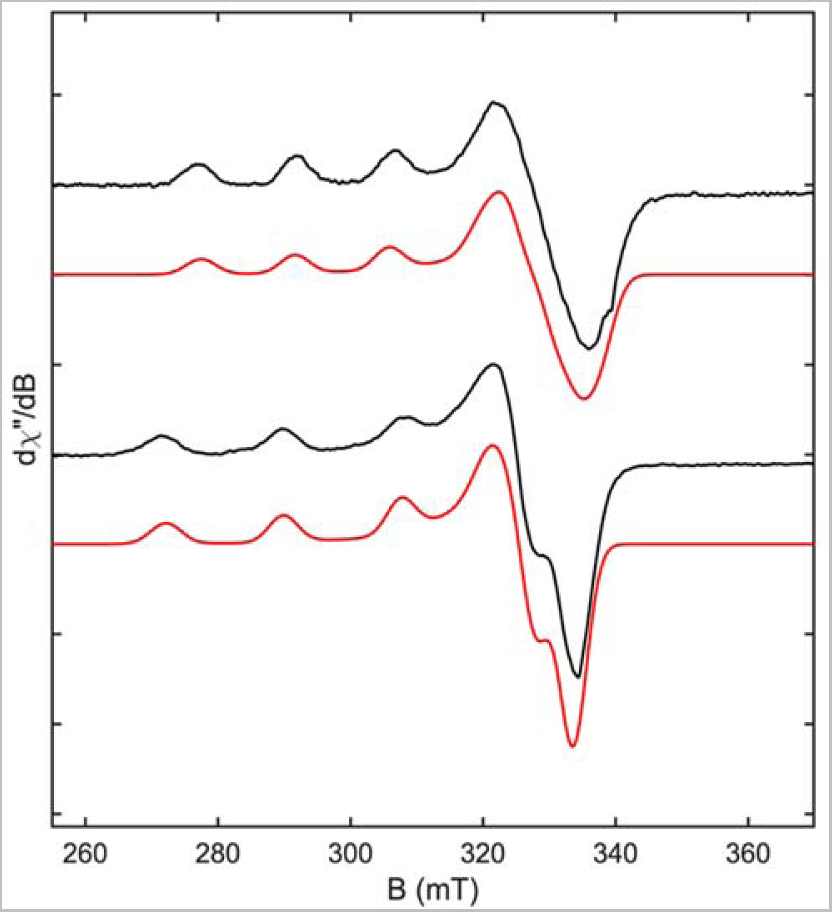
X-band EPR and simulated spectra of PMO^HG1^ and PMO^HG1^ H78A variant. The top black trace is the X-band CW spectrum from Cu(II) from PMO^HG1^. The top red trace is the simulated fit for WT PMO^HG1^ with the following parameters: g1 =2.245 g2 = 2.065 g3 = 2.005 A1 = 436 MHz gStrain1 = 0.03 gStrain1 = 0.03 gStrain1 = 0.03. The bottom black trace is the X-band CW spectrum from the Cu(II) bound to the PMO^HG1^ H78A variant. The lower red trace was simulated with the following parameters g1 2.245 g2 = 2.08 g3 = 2.05 A1 = 546 MHz gStrain1 = 0.03 gStrain1 = 0.035 gStrain1 = 0.02. All spectra were acquired at 40 K.

## References

1. Afzali, B., Lombardi, G., Lechler, R. I. (2008). Pathways of major histocompatibility complex allorecognition. Curr Op Organ Trans, 13(4), 438–444. https://doi.org/10.1097/MOT.0b013e328309ee31

2. Agger, J. W., Isaksen, T., Varnai, A., Vidal-Melgosa, S., Willats, W. G. T., Ludwig, R., … Westereng, B. (2014). Discovery of LPMO activity on hemicelluloses shows the importance of oxidative processes in plant cell wall degradation. Proc Natl Acad Sci USA, 111(17), 6287–6292. https://doi.org/10.1073/pnas.1323629111

3. Anna, S., Javier, P.-G., Mark, F., Louise, G. N. (2012). Physiological significance of network organization in fungi. Eukaryot Cell, 11(11), 1345–1352. https://doi.org/10.1128/EC.00213-12

4. Askarian, F., Uchiyama, S., Masson, H., Sørensen, H. V., Golten, O., Bunæs, A. C., … Vaaje-Kolstad, G. (2021). The lytic polysaccharide monooxygenase CbpD promotes *Pseudomonas aeruginosa* virulence in systemic infection. Nat Commun, 12(1), 1230. https://doi.org/10.1038/s41467-021-21473-0

5. Bastiaans, E., Debets, A. J. M., Aanen, D. K. (2016). Experimental evolution reveals that high relatedness protects multicellular cooperation from cheaters. Nat Commun, 7, 11435. https://doi.org/10.1038/ncomms11435

6. Bhowmick, R., Ghosal, A., Das, B., Koley, H., Saha, D. R., Ganguly, S., … Chatterjee, N. S. (2008). Intestinal adherence of *Vibrio cholerae* involves a coordinated interaction between colonization factor GbpA and mucin. Infection Immun, 76(11), 4968–4977. https://doi.org/10.1128/IAI.01615-07

7. Chiu, E., Hijnen, M., Bunker, R. D., Boudes, M., Rajendran, C., Aizel, K., … Coulibaly, F. (2015). Structural basis for the enhancement of virulence by viral spindles and their in vivo crystallization. Proc Natl Acad Sci, 112(13), 3973–3978. https://doi.org/10.1073/pnas.1418798112

8. Clark, J. (2003). Plasmodial incompatibility in the myxomycete *Didymium squamulosum*. Mycologia, 95(1), 24–26. https://doi.org/10.1080/15572536.2004.11833128

9. Corinne, C., Witold, D., A., T. E., Alexandra, G.-F., Léa, I., Benoît, P., … Asen, D. (2022). Fungal gasdermin-like proteins are controlled by proteolytic cleavage. Proc Natl Acad Sci, 119(7), e2109418119. https://doi.org/10.1073/pnas.2109418119

10. Courtade, G., Forsberg, Z., Heggset, E. B., Eijsink, V. G. H., Aachmann, F. L. (2018). The carbohydrate-binding module and linker of a modular lytic polysaccharide monooxygenase promote localized cellulose oxidation. J Biol Chem, 293(34), 13006–13015. https://doi.org/10.1074/jbc.RA118.004269

11. Couturier, M., Ladevèze, S., Sulzenbacher, G., Ciano, L., Fanuel, M., Moreau, C., … Berrin, J.-G. G. (2018). Lytic xylan oxidases from wood-decay fungi unlock biomass degradation. Nat Chem Biol, 14(3), 306–310.https://doi.org/10.1038/nchembio.2558

12. Danneels, B., Tanghe, M., Desmet, T. (2018). Structural features on the substrate-binding surface of fungal lytic polysaccharide monooxygenases determine their oxidative regioselectivity. Biotechnol J, https://doi.org/10.1002/biot.201800211

13. Daskalov, A., Gladieux, P., Heller, J., Glass, N. L. (2019). Programmed cell death in *Neurospora crassa* is controlled by the allorecognition determinant *rcd-1*. Genetics, 213(4), 1387–1400. https://doi.org/10.1534/genetics.119.302617

14. Daskalov, A., Glass, N. L. (2022). Gasdermin and gasdermin-like pore-forming proteins in invertebrates, fungi and bacteria. J Mol Biol, 434(4), 167273. https://doi.org//10.1016/j.jmb.2021.167273

15. Daskalov, A., Mitchell, P. S., Sandstrom, A., Vance, R. E., Glass, N. L. (2020). Molecular characterization of a fungal gasdermin-like protein. Proc Natl Acad Sci USA, 117(31), 18600LP – 18607. https://doi.org/10.1073/pnas.2004876117

16. De Tomaso, A. W., Nyholm, S. V, Palmeri, K. J., Ishizuka, K. J., Ludington, W. B., Mitchel, K., Weissman, I. L. (2005). Isolation and characterization of a protochordate histocompatibility locus. Nature, 438(7067), 454–459. https://doi.org/10.1038/nature04150

17. Debets, A. J. M., Dalstra, H. J. P., Slakhorst, M., Koopmanschap, B., Hoekstra, R. F., Saupe, S. J. (2012). High natural prevalence of a fungal prion. Proc Natl Acad Sci USA, 109(26), 10432LP – 10437. https://doi.org/10.1073/pnas.1205333109

18. Distel, D. L., Amin, M., Burgoyne, A., Linton, E., Mamangkey, G., Morrill, W., … Yang, J. (2011). Molecular phylogeny of *Pholadoidea Lamarck*, 1809 supports a single origin for xylotrophy (wood feeding) and xylotrophic bacterial endosymbiosis in Bivalvia. Mol Phylogen Evol, 61(2), 245–254. https://doi.org/10.1016/j.ympev.2011.05.019

19. Frandsen, K. E. H., Simmons, T. J., Dupree, P., Poulsen, J.-C. N., Hemsworth, G. R., Ciano, L., … Walton, P. H. (2016). The molecular basis of polysaccharide cleavage by lytic polysaccharide monooxygenases. Nat Chem Biol, 12(4), 298–303. https://doi.org/10.1038/nchembio.2029

20. Freitag, M., Hickey, P. C., Raju, N. B., Selker, E. U., Read, N. D. (2004). GFP as a tool to analyze the organization, dynamics and function of nuclei and microtubules in *Neurospora crassa*. Fungal Genet Biol, 41(10), 897–910. https://doi.org/10.1016/j.fgb.2004.06.008

21. Fu, C., Ao, J., Dettmann, A., Seiler, S., Free, S. J. (2014). Characterization of the *Neurospora crassa* cell fusion proteins, HAM-6, HAM-7, HAM-8, HAM-9, HAM-10, AMPH-1 and WHI-2. PLOS ONE, 9(10), e107773. https://doi.org/10.1371/journal.pone.0107773

22. Gerlt, J. A., Bouvier, J. T., Davidson, D. B., Imker, H. J., Sadkhin, B., Slater, D. R., Whalen, K. L. (2015). Enzyme function initiative-enzyme similarity tool (EFI-EST): A web tool for generating protein sequence similarity networks. Biochimica et Biophysica Acta https://doi.org/10.1016/j.bbapap.2015.04.015

23. Gibbs, A. K., Mark, U. L., Greenberg, E. P. (2008). Genetic determinants of self identity and social recognition in bacteria. Science, 321(5886), 256–259. https://doi.org/10.1126/science.1160033

24. Gibbs, K. A., Greenberg, E. P. (2011). Territoriality in Proteus: advertisement and aggression. Chemical Reviews, 111(1), 188–194. https://doi.org/10.1021/cr100051v

25. Gonçalves, A. P., Glass, N. L. (2020). Fungal social barriers: to fuse, or not to fuse, that is the question. Commun Integ Biol, 13(1), 39–42. https://doi.org/10.1080/19420889.2020.1740554

26. Gonçalves, A. P., Heller, J., Rico-Ramírez, A. M., Daskalov, A., Rosenfield, G., Glass, N. L. (2020). Conflict, competition, and cooperation regulate social interactions in filamentous fungi. Annu Rev Microbiol, 74, 693–712. https://doi.org/10.1146/annurev-micro-012420-080905

27. Gonçalves, A. P., Heller, J., Span, E. A. E., Rosenfield, G., Do, H. P., Palma-Guerrero, J., … Glass, N. L. (2019). Allorecognition upon fungal cell-cell contact determines social dooperation and impacts the acquisition of multicellularity. Curr Biol, 29(18), 1–12. https://doi.org/10.1016/j.cub.2019.07.060

28. Grum-Grzhimaylo, A. A., Bastiaans, E., van den Heuvel, J., Berenguer Millanes, C., Debets, A. J. M., Aanen, D. K. (2021). Somatic deficiency causes reproductive parasitism in a fungus. Nat Commun, 12(1), 783. https://doi.org/10.1038/s41467-021-21050-5

29. Gunawardena, N. D., Schrott, V., Richardson, C., Cole, T. N., Corey, C. G., Wang, Y., … Bullock, G. C. (2016). Aconitase: An iron sensing regulator of mitochondrial oxidative metabolism and erythropoiesis. Blood, 128(22), 74. https://doi.org/10.1182/blood.V128.22.74.74

30. Heller, J., Clavé, C., Gladieux, P., Saupe, S. J., Glass, N. L. (2018). NLR surveillance of essential SEC-9 SNARE proteins induces programmed cell death upon allorecognition in filamentous fungi. Proc Natl Acad Sci USA 115(10), E2292 LP-E2301. https://doi.org/10.1073/pnas.1719705115

31. Heller, J., Zhao, J., Rosenfield, G., Kowbel, D. J., Gladieux, P., Glass, N. L. (2016). Characterization of greenbeard genes involved in long-distance kind discrimination in a microbial eukaryote. PLoS Biol, 14(4), 1–29. https://doi.org/10.1371/journal.pbio.1002431

32. Hemsworth, G. R., Henrissat, B., Davies, G. J., Walton, P. H. (2014). Discovery and characterization of a new family of lytic polysaccharide monooxygenases. Nat Chem Biol, 10(2), 122–126. https://doi.org/10.1038/nchembio.1417

33. Hüttner, S., Várnai, A., Petrović, D. M., Bach, C. X., Kim Anh, D. T., Thanh, V. N., … Olsson, L. (2019). Specific xylan activity revealed for AA9 lytic polysaccharide monooxygenases of the thermophilic fungus *Malbranchea cinnamomea* by functional characterization. Appl Environ Microbiol, 85(23). https://doi.org/10.1128/AEM.01408-19

34. Jeffery, C. J. (1999). Moonlighting proteins. Trends Biochem Sci, 24(1), 8–11. https://doi.org//10.1016/S0968-0004(98)01335-8

35. Jeffery, C. J. (2018). Protein moonlighting: what is it, and why is it important? Phil Trans Royal Soc B: Biol Sci, 373(1738), 20160523. https://doi.org/10.1098/rstb.2016.0523

36. Johansen, K. S. (2016). Discovery and industrial applications of lytic polysaccharide mono-oxygenases. Biochem Soc Trans, 44(1), 143–149. http://www.biochemsoctrans.org/content/44/1/143.abstract

37. Kittl, R., Kracher, D., Burgstaller, D., Haltrich, D., Ludwig, R. (2012). Production of four *Neurospora crassa* lytic polysaccharide monooxygenases in *Pichia pastoris* monitored by a fluorimetric assay. Biotechnol Biofuels, 5(1), 79–92. https://doi.org/10.1186/1754-6834-5-79

38. Kruer-Zerhusen, N., Alahuhta, M., Lunin, V. V., Himmel, M. E., Bomble, Y. J., Wilson, D. B. (2017). Structure of a *Thermobifida fusca* lytic polysaccharide monooxygenase and mutagenesis of key residues. Biotechnol Biofuels, 10(1), 243. https://doi.org/10.1186/s13068-017-0925-7

39. Kundert, P., Shaulsky, G. (2019). Cellular allorecognition and its roles in Dictyostelium development and social evolution. Internatl J Dev Biol, 63(8-9–10), 383–393. https://doi.org/10.1387/ijdb.190239gs

40. Kuzdzal-Fick J., J., Fox A., S., Strassmann E., J., Queller C., D. (2011). High relatedness is necessary and sufficient to maintain multicellularity in Dictyostelium. Science, 334(6062), 1548–1551. https://doi.org/10.1126/science.1213272

41. Laurent, C. V. F. P., Breslmayr, E., Tunega, D., Ludwig, R., Oostenbrink, C. (2019). Interaction between cellobiose cehydrogenase and lytic polysaccharide monooxygenase. Biochem. https://doi.org/10.1021/acs.biochem.8b01178

42. Levasseur, A., Drula, E., Lombard, V., Coutinho, P. M., Henrissat, B. (2013). Expansion of the enzymatic repertoire of the CAZy database to integrate auxiliary redox enzymes. Biotechnol Biofuels, 6(1). https://doi.org/10.1186/1754-6834-6-41

43. Liu, B., Kognole, A. A., Wu, M., Westereng, B., Crowley, M. F., Kim, S., … Sandgren, M. (2018). Structural and molecular dynamics studies of a C1-oxidizing lytic polysaccharide monooxygenase from *Heterobasidion irregulare* reveal amino acids important for substrate recognition. FEBS J, 285(12), 2225–2242. https://doi.org/10.1111/febs.14472

44. Loose, J. S. M. M., Forsberg, Z., Fraaije, M. W., Eijsink, V. G. H. H., Vaaje-Kolstad, G. (2014). A rapid quantitative activity assay shows that the *Vibrio cholerae* colonization factor GbpA is an active lytic polysaccharide monooxygenase. FEBS Letts, 588(18), 3435–3440. https://doi.org/10.1016/j.febslet.2014.07.036

45. M., J. S., J., T. W., K., M. K., Bradley, K., I., S. E. (2020). Kinetic analysis of amino acid radicals formed in H_2_O_2_-driven CuI LPMO reoxidation implicates dominant homolytic reactivity. Proc Natl Acad Sci USA, 117(22), 11916–11922. https://doi.org/10.1073/pnas.1922499117

46. Maddi, A., Bowman, S. M., Free, S. J. (2009). Trifluoromethanesulfonic acid-based proteomic analysis of cell wall and secreted proteins of the ascomycetous fungi *Neurospora crassa* and *Candida albicans*. Fungal Genet Biol. https://doi.org/10.1016/j.fgb.2009.06.005

47. Maddi, A., Dettman, A., Fu, C., Seiler, S., Free, S. J. (2012). WSC-1 and HAM-7 are MAK-1 MAP kinase pathway sensors required for cell wall integrity and hyphal fusion in *Neurospora crassa*. PLoS ONE, 7(8). https://doi.org/10.1371/journal.pone.0042374

48. Margolin, B. S., Freitag, M., Selker, E. U. (1997). Improved plasmids for gene targeting at the his-3 locus of *Neurospora crassa* by electroporation. Fungal Genet Reports, 44(1), 34–36. https://doi.org/10.4148/1941-4765.1281

49. Marino, J., Paster, J., Benichou, G. (2016). Allorecognition by T lymphocytes and allograft rejection. Front Immunol. https://www.frontiersin.org/article/10.3389/fimmu.2016.00582

50. McCluskey, K., Wiest, A., Plamann, M. (2010). The Fungal Genetics Stock Center: a repository for 50 years of fungal genetics research. J Biosci, 35(1), 119–126. https://doi.org/10.1007/s12038-010-0014-6

51. Monclaro, A. V, Petrović, D. M., Alves, G. S. C., Costa, M. M. C., Midorikawa, G. E. O., Miller, R. N. G., … Várnai, A. (2020). Characterization of two family AA9 LPMOs from *Aspergillus tamarii* with distinct activities on xyloglucan reveals structural differences linked to cleavage specificity. PLOS ONE, 15(7), e0235642. https://doi.org/10.1371/journal.pone.0235642

52. Moreau, C., Tapin-Lingua, S., Grisel, S., Gimbert, I., Le Gall, S., Meyer, V., … Villares, A. (2019). Lytic polysaccharide monooxygenases (LPMOs) facilitate cellulose nanofibrils production. Biotechnol Biofuels, 12(1), 156. https://doi.org/10.1186/s13068-019-1501-0

53. Palma-Guerrero, J., Hall, C. R., Kowbel, D., Welch, J., Taylor, J. W., Brem, R. B., Glass, N. L. (2013). Genome wide association identifies novel loci involved in fungal communication. PLOS Genetics, 9(8), e1003669. https://doi.org/10.1371/journal.pgen.1003669

54. Pandey, K. K. (1960). Incompability in Abutilon “Hybridum.” Amer J Bot, 47(10), 877– 883. https://doi.org//10.1002/j.1537-2197.1960.tb07180.x

55. Phillips, C. M., Beeson, W. T., Cate, J. H., Marletta, M. A. (2011). Cellobiose dehydrogenase and a copper-dependent polysaccharide monooxygenase potentiate cellulose degradation by *Neurospora crassa*. ACS Chem Biol, 6(12), 1399–1406. https://doi.org/10.1021/cb200351y

56. Piatigorsky, J. (1998). Multifunctional lens crystallins and corneal enzymes. More than meets the eye. Annal New York Acad Sci, 842, 7–15. https://doi.org/10.1111/j.1749-6632.1998.tb09626.x

57. Pinheiro, G. L., Correa, R. F., Cunha, R. S., Cardoso, A. M., Chaia, C., Clementino, M. M., … Frasés, S. (2015). Isolation of aerobic cultivable cellulolytic bacteria from different regions of the gastrointestinal tract of giant land snail *Achatina fulica*. Front Microbiol, 6, 860. https://doi.org/10.3389/fmicb.2015.00860

58. Rico-Ramírez, A. M., Gonçalves, A. P., Louise Glass, N. (2022). Fungal cell death: The beginning of the end. Fungal Genet Biol, 159, 103671. https://doi.org//10.1016/j.fgb.2022.103671

59. Rieder, L., Petrović, D., Väljamäe, P., Eijsink, V. G. H., Sørlie, M. (2021). Kinetic characterization of a putatively chitin-active LPMO reveals a preference for soluble substrates and absence of monooxygenase activity. ACS Catalysis, 11(18), 11685– 11695. https://doi.org/10.1021/acscatal.1c03344

60. Rosengarten, R. D., Nicotra, M. L. (2011). Model systems of invertebrate allorecognition. Curr Biol, 21(2), R82–R92. https://doi.org/10.1016/j.cub.2010.11.061

61. Sabbadin, F., Hemsworth, G. R., Ciano, L., Henrissat, B., Dupree, P., Tryfona, T., … McQueen-Mason, S. J. (2018). An ancient family of lytic polysaccharide monooxygenases with roles in arthropod development and biomass digestion. Nat Commun, 9(1). https://doi.org/10.1038/s41467-018-03142-x

62. Saupe, S. J. (2011). The [Het-s] prion of *Podospora anserina* and its role in heterokaryon incompatibility. Sem Cell Dev Biol, 22(5), 460–468. https://doi.org//10.1016/j.semcdb.2011.02.019

63. Saupe, S. J. (2020). Amyloid signaling in filamentous fungi and bacteria. Annu Rev Microbiol, 74(1), 673–691. https://doi.org/10.1146/annurev-micro-011320-013555

64. Schneider, C. A., Rasband, W. S., Eliceiri, K. W. (2012). NIH Image to ImageJ: 25 years of image analysis. Nat Meth, 9(7), 671–675. https://doi.org/10.1038/nmeth.2089

65. Shaulsky, G., Kessin, R. H. (2007). The cold war of the social amoebae. Curr Biol, 17(16), R684–R692. https://doi.org/10.1016/j.cub.2007.06.024

66. Span, E. A., Suess, D. L. M., Deller, M. C., Britt, R. D., Marletta, M. A. (2017). The role of the secondary coordination sphere in a fungal polysaccharide monooxygenase. ACS Chem Biol, 12(4), 1095–1103. https://doi.org/10.1021/acschembio.7b00016

67. Støpamo, F. G., Røhr, Å. K., Mekasha, S., Petrović, D. M., Várnai, A., Eijsink, V. G. H. (2021). Characterization of a lytic polysaccharide monooxygenase from *Aspergillus fumigatus* shows functional variation among family AA11 fungal LPMOs. J Biol Chem, 297(6). https://doi.org/10.1016/j.jbc.2021.101421

68. Tan, T. C., Kracher, D., Gandini, R., Sygmund, C., Kittl, R., Haltrich, D., … Divne, C. (2015). Structural basis for cellobiose dehydrogenase action during oxidative cellulose degradation. Nat Commun, 6(May). https://doi.org/10.1038/ncomms8542

69. Vaaje-Kolstad, G., Westereng, B., Horn, S. J., Liu, Z., Zhai, H., Sørlie, M., Eijsink, V. G. H. (2010). An oxidative enzyme boosting the enzymatic conversion of recalcitrant polysaccharides. Science, 330(6001), 219–222. http://science.sciencemag.org/content/330/6001/219.abstract

70. Vogel, H. L. (1956). A convenient medium for *Neurospora*. Microbial Genetics Bulletin, 13, 42–43.

71. Vu, V. V, Beeson, W. T., Phillips, C. M., Cate, J. H. D., Marletta, M. A. (2014). Determinants of regioselective hydroxylation in the fungal polysaccharide monooxygenases. J Amer Chem Soc, 136(2), 562–565. https://doi.org/10.1021/ja409384b

72. Vu, V. V, Hangasky, J. A., Detomasi, T. C., Henry, S. J. W., Ngo, S. T., Span, E. A., Marletta, M. A. (2019). Substrate selectivity in starch polysaccharide monooxygenases. J Biol Chem, 294(32), 12157–12166. https://doi.org//10.1074/jbc.RA119.009509

73. Vu, V. V, Marletta, M. A. (2016). Starch-degrading polysaccharide monooxygenases. Cell Mol Life Sci. https://doi.org/10.1007/s00018-016-2251-9

74. Wang, D., Li, J., Salazar-Alvarez, G., McKee, L. S., Srivastava, V., Sellberg, J. A., … Hsieh, Y. S. Y. (2018). Production of functionalised chitins assisted by fungal lytic polysaccharide monooxygenase. Green Chem, 20(9), 2091–2100. https://doi.org/10.1039/C8GC00422F

75. Wang, D., Li, J., Wong, A. C. Y., Aachmann, F. L., Hsieh, Y. S. Y. (2018). A colorimetric assay to rapidly determine the activities of lytic polysaccharide monooxygenases. Biotechnol Biofuels. https://doi.org/10.1186/s13068-018-1211-z

76. Westergaard, M., Mitchell, H. K. (1947). *Neurospora* V. A synthetic medium favoring sexual reproduction. Amer J Bot, 34(10), 573–577. https://doi.org/10.1002/j.1537-2197.1947.tb13032.x

77. Wojtowicz, W. M., Flanagan, J. J., Millard, S. S., Zipursky, S. L., Clemens, J. C. (2004). Alternative splicing of *Drosophila* Dscam generates axon guidance receptors that exhibit isoform-specific homophilic binding. Cell, 118(5), 619–633. https://doi.org//10.1016/j.cell.2004.08.021

78. Wojtowicz, W. M., Wu, W., Andre, I., Qian, B., Baker, D., Zipursky, S. L. (2007). A vast repertoire of Dscam binding specificities arises from modular interactions of variable Ig domains. Cell, 130(6), 1134–1145. https://doi.org//10.1016/j.cell.2007.08.026

79. Wood, T. M. B. T.-M. in E. (1988). Preparation of crystalline, amorphous, and dyed cellulase substrates. In Biomass Part A: Cellulose and Hemicellulose (Vol. 160, pp. 19–25). Academic Press. https://doi.org//10.1016/0076-6879(88)60103-0

80. Wu, V. W., Thieme, N., Huberman, L. B., Dietschmann, A., Kowbel, D. J., Lee, J., … Glass, N. L. (2020). The regulatory and transcriptional landscape associated with carbon utilization in a filamentous fungus. Proc Natl Acad Sci USA, 117(11), 6003– 6013. https://doi.org/10.1073/pnas.1915611117

81. Yoshito, H., Yuhei, T., Masahiko, S., Takako, S., Lixy, Y., Hisaaki, T., … Hitoshi, S. (2008). Mechanism of self-sterility in a hermaphroditic chordate. Science, 320(5875), 548–550. https://doi.org/10.1126/science.1152488

82. Zhang, D.-X., Nuss, D. L. (2016). Engineering super mycovirus donor strains of chestnut blight fungus by systematic disruption of multilocus genes. Proc Natl Acad Sci USA, 113(8), 2062–2067. https://doi.org/10.1073/pnas.1522219113

83. Zheng, W., Francesc, L.-G., Nina, L., Marta, F., Ralph, C., Frances, T., P., T. J. (2014). Global gene expression and focused knockout analysis reveals genes associated with fungal fruiting body development in *Neurospora crassa*. Eukaryot Cell, 13(1), 154–169. https://doi.org/10.1128/EC.00248-13

